# Evaluating microbial network inference methods: Moving beyond synthetic data with reproducibility-driven benchmarks

**DOI:** 10.1101/2025.07.05.663212

**Authors:** Zahra Ghaeli, Rosa Aghdam, Changiz Eslahchi

## Abstract

Microbial network inference is an essential approach for revealing complex interactions within microbial communities. However, the lack of experimentally validated gold standards presents a significant obstacle in evaluating the accuracy and biological relevance of inferred networks. This study delivers a comprehensive comparative assessment of six widely used microbial network inference algorithms—gCoda, OIPCQ, S-E(glasso), S-E(mb), SPRING, and SparCC—using four diverse real-world microbiome datasets alongside multiple types of generated data, including synthetic, noisy, and bootstrap-derived datasets. Our evaluation framework extends beyond conventional synthetic benchmarking by emphasizing reproducibility-focused assessments grounded in biologically realistic perturbations. We show that bootstrap resampling and low-level noisy datasets (≤10% perturbation) effectively preserve key statistical properties of real microbiome data, such as diversity indices, abundance distributions, and sparsity patterns. Conversely, synthetic datasets generated via the widely used SPIEC-EASI method exhibit substantial divergence from real data across these metrics. We find that while SparCC demonstrates superior robustness across varying data conditions, other methods tend to produce inflated performance metrics when evaluated on unrealistic synthetic networks. Notably, several algorithms fail to distinguish between structured and random networks, highlighting issues of structural insensitivity and the limitations of overreliance on synthetic benchmarks. To address these challenges, we propose a reproducibility-centered benchmarking framework that prioritizes real-data-derived perturbations and mandates rigorous statistical validation of synthetic datasets before their use. This work provides critical insights and practical guidance for the microbiome research community, aiming to foster more reliable and ecologically meaningful microbial network inference in the absence of a true ground truth.

## 1 Introduction

The term microbiota refers to a community of microorganisms—including bacteria, archaea, protists, fungi, and viruses—that coexist within a shared environment. In contrast, the term microbiome includes not only these microorganisms but also their genetic material and the environmental context in which they reside [1]. With advances in sequencing technologies, comprehensive human microbiome datasets have become available, enabling extensive research into the microbiome’s role in human health and disease [2–4]. The microbiome plays a critical role in the development, progression, and treatment of various diseases, including cancer [5]. Dysbiosis—an imbalance in the microbial community—has been linked to colorectal, breast, lung, and prostate cancers [6], often through its impact on metabolic, inflammatory, and immunological pathways. The gut microbiome is also implicated in neurodegenerative disorders such as Alzheimer’s disease, where it influences both metabolite production and genetic factors involved in disease progression [7]. Moreover, a bidirectional relationship has been observed between the gut microbiome and schizophrenia: not only does microbial composition influence the disorder’s onset and progression, but the disease itself alters the microbiome structure [8]. Microbial interactions are often classified broadly as positive, neutral, or negative, reflecting patterns such as co-occurrence, independence, or mutual exclusion among operational taxonomic units (OTUs) [9].

In microbiome studies, 16S rRNA gene sequencing and whole-genome sequencing are commonly used to group sequences sharing more than 97% similarity into OTUs. The resulting data are typically structured as abundance matrices, where rows represent samples and columns represent OTUs. These matrices often contain many zeros—due either to true biological absence or technical limitations such as insufficient sequencing depth—making interpretation difficult [10, 11]. Furthermore, due to variability in sequencing depth, microbiome data are inherently compositional—that is, they represent relative rather than absolute abundances. This constant-sum constraint introduces dependencies among taxa, potentially leading to spurious correlations, particularly negative ones. To address these issues, log-ratio transformations—such as the centered log-ratio (CLR) method—are commonly employed, as they allow more valid statistical analyses by accounting for the compositional nature of the data [12, 13].

Inferring microbial interaction networks is a key task in microbiome research, as these networks provide insights into how different OTUs coexist, compete, and influence host health [9, 14, 15]. In such networks, nodes represent OTUs and edges denote potential biological associations, such as co-occurrence, competition, or mutual exclusion. These interactions are often inferred from co-occurrence patterns and occasionally supported by experimental validation [16]. Microbial networks have practical applications in identifying keystone OTUs, understanding community resilience, and guiding microbiome-targeted therapeutic strategies [17].

To reconstruct these networks from sequencing data, various computational algorithms have been developed, typically categorized into correlation-based and conditional dependence-based approaches. Early correlation-based methods include Pearson and Spearman correlations, followed by SparCC (Sparse Correlations for Compositional data) [18], which adjusts for compositional effects and assumes sparsity in the network. This line of work has led to more advanced methods such as CCLasso (Correlation inference for Compositional data through Lasso) [19] and REBACCA (regularized estimation of the basis covariance based on compositional data) [20], which further refine network inference under compositional constraints. Conditional dependence-based methods aim to capture direct associations between OTUs by accounting for indirect interactions. Prominent examples include the SPIEC-EASI (SParse InversE Covariance Estimation for Ecological Association Inference) [13], which uses graphical models adapted for compositional data, and SPRING (Semi-Parametric Rank-based Inference in Graphical models) algorithm [21], which employs partial correlations. More recent tools—such as CONet [22], meta-network [23], HARMONIES (Hybrid Approach foR MicrobiOme Network Inferences via Exploiting Sparsity) [24], FLASHWeave [25], and MicroNet-MIMRF (Microbial Network based on Mutual Information and Markov Random Fields) [26]—aim to improve robustness to sparsity, compositionality, and technical noise in OTU-level microbiome data.

One of the key challenges in microbial network inference is the absence of a validated reference for real microbial interactions. As a result, many studies rely on synthetic datasets generated from artificial topologies (e.g., cluster, band, scale-free networks) to evaluate algorithm performance [13, 21, 25–27]. However, such artificial networks often fail to capture the ecological complexity, nonlinearity, and context-dependency of real microbial ecosystems [9, 16]. In real communities, microbial interactions are shaped by a variety of factors, including metabolic dependencies, environmental conditions, and host responses—features that are difficult to mimic in idealized graph structures. This disconnect raises concerns about the biological relevance of results obtained from synthetic evaluations and underscores the need for alternative validation strategies grounded in real-world data.

In this paper, we address the critical challenge of evaluating microbial network inference algorithms by conducting an extensive comparison of six widely adopted methods: gCoda [27] a graphical model tailored for compositional data, based on a normal-logistic distribution and sparse inverse covariance estimation, OIPCQ (Order Independent PC-based algorithm using Conditional Quantification), initially developed for gene regulatory networks [28] extracts direct dependencies between microorganisms based on the concept of conditional mutual information, S-E(glasso) (SPIAC-EASI with Graphical Lasso) [13] which is a part of the SPIEC-EASI framework, S-E(mb) (SPIEC-EASI with Meinshausen–Bühlmann neighborhood selection) [13], SPRING (Semi-Parametric Rank-based Inference in Graphical models) [21] a semi-parametric rank based method that estimates partial correlations while accounting for compositionality, and SparCC (Sparse Correlations for Compositional data) [18] a correlation-based method that infers robust associations by correcting for the compositional nature of the data using log-ratio transformations. We evaluated these algorithms on four real-world microbiome datasets to rigorously test their performance. Importantly, unlike traditional evaluations that primarily utilize synthetic data generated from predefined, idealized network topologies, our approach includes more statistically grounded and reproducibility-oriented benchmarks, such as bootstrap sampling and controlled noisy perturbations derived directly from real datasets. Our evaluation framework provides deeper insight into the reliability of network inference methods by systematically comparing performance across synthetic, noisy, and bootstrap-generated datasets. We highlight significant limitations in traditional synthetic data benchmarks, demonstrating their inadequacy in reflecting the genuine statistical complexity and structural properties of real microbiome datasets. Our results strongly advocate for reproducibility-focused benchmarks using real-data-derived perturbations to accurately assess methodological robustness. This study thus offers a clear and practical pathway for future methodological evaluations, emphasizing the critical importance of realistic data scenarios to support meaningful interpretation and robust performance assessment.

## 2 Materials and methods

### 2.1 Dataset sources

We utilized four publicly available microbiome datasets in this study. The first two, `amgut1` and `amgut2`, are provided in the `SpiecEasi` R package and correspond to two phases of the American Gut Project (AGP), a large-scale citizen science initiative studying the human gut microbiome. Both datasets are based on stool samples: `amgut1` includes 289 samples and 127 OTUs, while `amgut2` contains 296 samples and 138 OTUs [13]. The third dataset, denoted as `GUT`, was collected to examine the relationship between gut microbiome composition and susceptibility to *Plasmodium falciparum* infection [29]. It includes 195 stool samples and 128 OTUs [30]. The fourth dataset, referred to as `MOMS-PI`, originates from the NIH-funded Multi-Omic Microbiome Study: Pregnancy Initiative (MOMS-PI), which investigates the role of the microbiome in maternal and infant health. The dataset used in this study consists of 225 vaginal samples and 90 OTUs [31].

### 2.2 Data generation methods

To evaluate microbial network inference algorithms under varied conditions, we generated three types of datasets derived from the real data: synthetic data, noisy data, and bootstrap data.

#### Synthetic data

The SpiecEasi and SPRING methods generates synthetic data that mimics real data through the NORmal To Anything (NORTA) algorithm [32]. Both methods simulate count-like data by taking as input a predefined network topology and a real dataset, and producing synthetic samples that reflect the structure of the input network. We used a total of 11 input network topologies for data generation. These included three predefined structures—scale-free, banded, and clustered graphs (from `SpiecEasi` R package)—and two random networks generated using the `Erdos-Renyi` function from the `igraph` R package [33], as well as six networks inferred from real data using the gCoda, OIPCQ, S-E(glasso), S-E(mb), SPRING, and SparCC algorithms. For each of the 11 topologies, we generated synthetic datasets using both the SPIEC-EASI (SE) and SPRING (SP) methods, resulting in a total of 22 synthetic datasets for each real dataset (11 topologies *×* 2 methods).

#### Noisy data

To evaluate the sensitivity of network inference methods to small and big perturbations in the input data, we generated noisy datasets by introducing controlled random noise to real microbiome data. For each OTU (i.e., column of the abundance matrix), we computed its mean and variance. Then, for a specified noise level (5%, 10%, 20%, 30%, 40%, or 50%), we randomly selected that percentage of entries within each column and add them with values drawn from a truncated normal distribution matching the original column’s mean and variance. This operation was performed independently for each OTU. The modified values were rounded to the nearest integer to maintain the count format of microbiome data.

#### Bootstrap data

To evaluate the reproducibility of network inference in the presence of sample-level variability, we generated bootstrap datasets by resampling complete samples (i.e., rows) from each original dataset with replacement. Each bootstrap replicate contained the same number of samples as the original, thus maintaining the overall OTU distribution while introducing stochastic variation due to sampling. This approach simulates the scenario of analyzing a similar cohort with comparable characteristics but slight differences in sample composition.

Thus, using three types of datasets derived from the real data, synthetic, noisy, and bootstrap, we generated a total of 29 distinct dataset types for each real dataset: 22 synthetic datasets (from 11 network topologies using two generation methods), 6 noisy datasets at varying noise levels, and 1 set of bootstrap replicates. To assess algorithm robustness, both the synthetic and noisy datasets were generated 100 times. The number of bootstrap replicates matched the number of samples in the original dataset.

### 2.3 Comparing generated data with real data

It is crucial that the generated datasets, synthetic, noisy, and bootstrap, resemble the real microbiome data in meaningful ways, as this similarity ensures that downstream evaluations of network inference methods are both realistic and reliable. To assess the faithfulness of the generated data, we conducted three types of comparisons: diversity analysis, matrix entry similarity, and distributional alignment.

#### Diversity analysis

We computed three alpha diversity indices for each sample in both the real and generated datasets: Richness, the Shannon index, and the inverse Simpson index [34]. These metrics capture different aspects of microbial community diversity, including species richness and evenness. The Richness refers to the number of species present in a sample. The Shannon index is defined as:

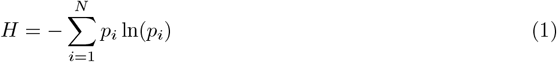

where *N* is the number of OTUs in a sample, and *p*_*i*_ is the probability of observing the *i*-th OTU, estimated by its relative abundance (i.e., the proportion of total counts in the sample). The Simpson index is calculated as:

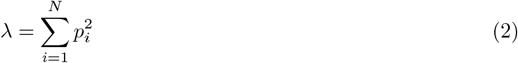

and the Inverse Simpson index is given by 1*/λ*. We computed these indices using the `diversity` function from the `vegan` R package [35]. To statistically compare the distributions of diversity scores between real and generated data, we applied the Wilcoxon rank-sum test. Since each synthetic and noisy dataset was generated 100 times, p-values were calculated for each replicate to evaluate variability and statistical significance. However, examining diversity indices alone is not sufficiently precise, and one cannot conclude that two datasets lack a significant difference solely based on the absence of a statistically significant difference in their indices. It is possible that synthetic data may not show a statistically significant difference in a particular index, yet in reality, they may differ substantially due to variations in the matrix entries, thus masking the absence of a significant difference. Therefore, since we are comparing real data with synthetic, noisy, and bootstrap data—not healthy versus diseased data—it is important that the results of diversity indices be interpreted in a way that considers the degree of agreement between the data entries and the real data.

#### Matrix entry similarity (F-score)

To evaluate the presence–absence agreement (binary overlap) between real and generated abundance matrices, we binarized all datasets by setting non-zero values to 1 and zeros to 0. For each generated matrix, we computed the number of true positives (TP), true negatives (TN), false positives (FP), and false negatives (FN) by comparing to the binarized real data. Using these values, we calculated the 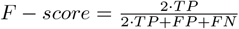 for each replicate to assess how closely the structure of presence/absence entries in the generated data matched the real data.

#### Distribution comparison (KS test)

To assess how well the distribution of OTU abundances was preserved in the generated datasets, we applied the Kolmogorov–Smirnov (KS) test. This non-parametric test evaluates whether two samples are drawn from the same distribution by comparing their empirical cumulative distribution functions (ECDFs). The KS statistic is defined as:

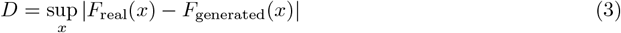

where *F*_real_(*x*) and *F*_generated_(*x*) are the empirical cumulative distribution functions of the real and generated OTU abundance values, respectively, and sup_*x*_ denotes the supremum over all values of *x*. The KS test was performed independently for each OTU, comparing its abundance distribution in the real dataset to that in each replicate of the generated dataset. We report the percentage of OTUs for which a significant difference was detected, based on the KS test at a standard significance threshold (*p* − *value <* .05).

### 2.4 Evaluation criteria for network comparison

To evaluate the performance of the six microbial network inference algorithms described above, we compared the inferred networks to reference networks using F-score metric. We adjusted the parameters of the methods such that the number of edges in the generated networks would fall within a range, for a fair comparison. The type of reference network varied depending on the data source:

For each synthetic dataset generated by the SE and SP methods, we used the underlying topology used (11 different topologies) for data generation as the reference. Each algorithm inferred a network from the synthetic data, and this inferred network was compared to the original input topology.

For noisy and bootstrap datasets, we used the network inferred from the corresponding real dataset as the reference (i.e., gold standard). Specifically, for evaluating a given method *A*, we first inferred a network from the real dataset using method *A*, and this resulting network served as the reference to evaluate method *A*’s performance on the corresponding noisy or bootstrap data. This ensures that each algorithm is evaluated consistently with respect to its own assumptions and inference framework. Each algorithm was applied to both the real and generated datasets, and the resulting networks were compared to assess reproducibility. In all cases, networks were represented as adjacency matrices, and similarity was computed using Precision, Recall and F-score which are define as follows:

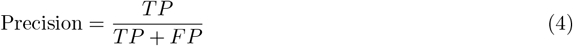

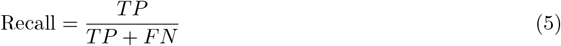

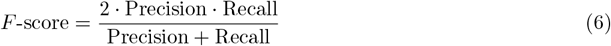

Here, TP is the number of edges present in both the inferred and reference networks, TN is the number of absent edges in both, FP are edges present only in the inferred network, and FN are edges present only in the reference network. This evaluation allows us to quantify both the accuracy of network inference on synthetic datasets and the stability of algorithmic output across noisy and bootstrap replicates. The overall workflow of the study is summarized in Figure 1. The diagram outlines the key stages, including data generation (synthetic, noisy, and bootstrap) and microbial network reconstruction, aimed at establishing a robust framework for evaluating new microbial network inference algorithms.

**Figure 1:**
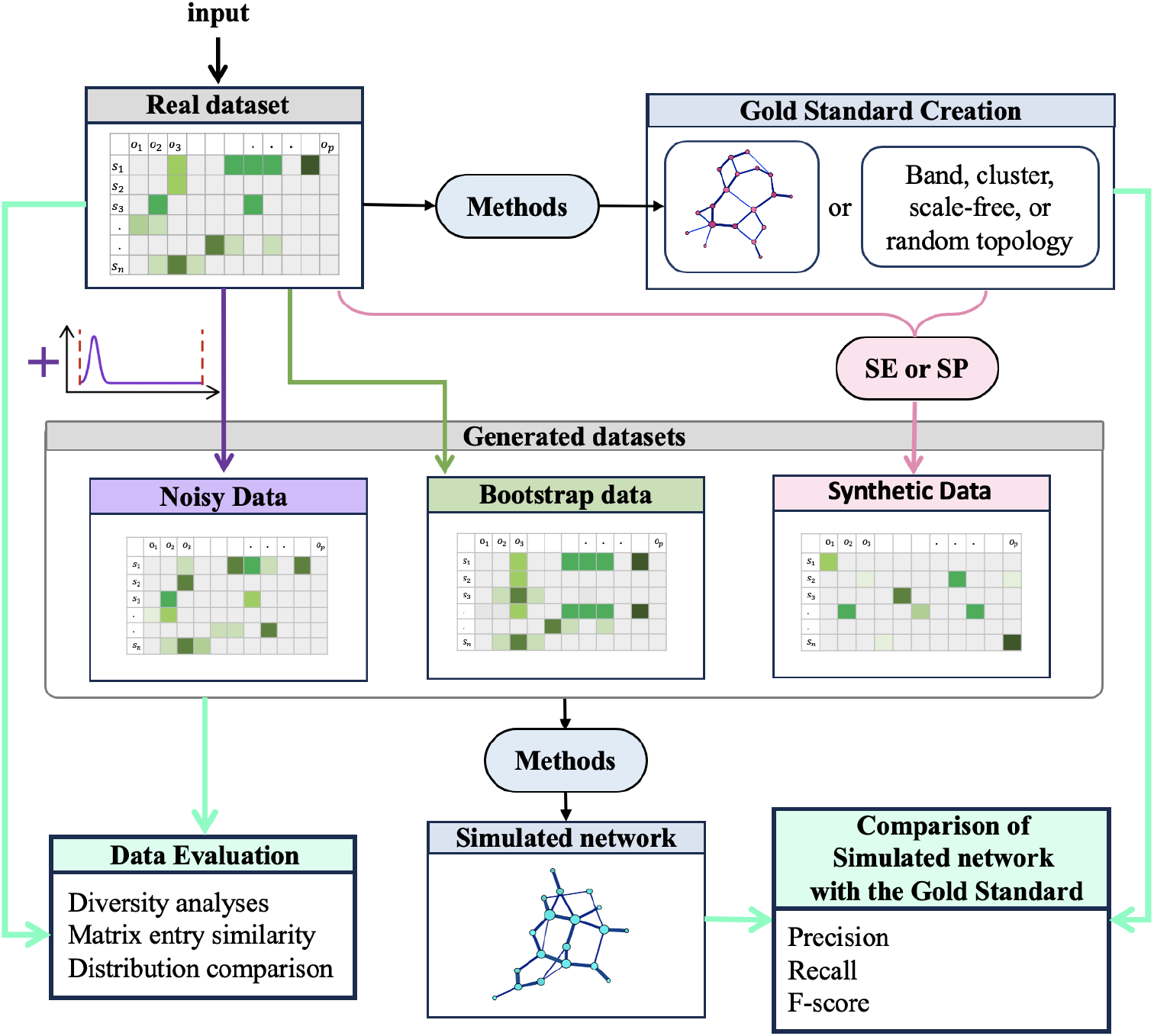
Overview of the study workflow, illustrating real data selection, generation of synthetic (SE or SP), noisy, and bootstrap datasets, data evaluation and comparative evaluation of microbial network inference algorithms. Methods: gCoda, OIPCQ, S-E(glasso), S-E(mb), SPRING, and SparCC.

## 3 Results

### 3.1 Comparison between real and generated data

We evaluated the similarity between the generated datasets (synthetic, noisy, and bootstrap) and the real microbiome datasets using three complementary criteria: diversity indices, matrix entry similarity (F-score), and distributional alignment via the KS test.

#### Diversity indices

The Wilcoxon rank-sum test was applied to compare the Richness, Shannon and Inverse Simpson index between real and generated samples across all datasets. As shown in Table 2, synthetic data generated by the SE method consistently exhibited statistically significant differences (*p* − *value <* 0.05) from the real data in all four datasets. The SP method also showed significant differences for `amgut1` and `amgut2`, but not for `GUT` and `MOMS-PI`. Noisy datasets with up to 10% perturbation did not significantly differ from the real data in terms of the Shannon index. However, for `GUT` and `MOMS-PI`, even 20% noise preserved similarity, while differences became significant for noise levels of 30% and higher. Bootstrap datasets showed no significant difference across all four datasets. The violin plots for the `amgut1`, `amgut2`, `GUT`, and (MOMS-PI) datasets, are shown in Figures S1 to S4. The results for the Inverse Simpson index are identical to those for the Shannon index (see Figures S5 to S8). Furthermore, for the Richness index, data generated by the SE method shows a statistically significant difference across all four real-world datasets. The SP method also shows a statistically significant difference for `amgut1` and `amgut2`, but not for `GUT` and `MOMS-PI` (see Figures S9 to S12). No statistically significant difference was observed between the noisy and Bootstrapped data and the real-world datasets.

**Table 1:** Summary of microbiome datasets used in this study, including sample size, number of OTUs, and data source.

| Dataset | Samples | OTUs | Source |
| --- | --- | --- | --- |
| `amgut1` | 289 | 127 | `SpiecEasi` R package [13] |
| `amgut2` | 296 | 138 | `SpiecEasi` R package [13] |
| `GUT` | 195 | 128 | https://github.com/sahatava/MixMPLN/blob/master/data/real_data.csv [30] |
| `MOMS-PI` | 225 | 90 | https://github.com/Nathan-Osborne/SINC/blob/master/DataApplication/OTU.csv [31] |

**Table 2:** Results of the Wilcoxon rank-sum test comparing the Shannon diversity index between real and generated datasets. A red cross (×) indicates statistically significant differences (*p* − *value <* .05).

|  | amgut1 | amgut2 | GUT | MOMS-PI |
| --- | --- | --- | --- | --- |
| Synthetic data SE | × | × | × | × |
| Synthetic data SP | × | × |  |  |
| Bootstrap |  |  |  |  |
| Noisy .05 |  |  |  |  |
| Noisy .1 |  |  |  |  |
| Noisy .2 | × | × |  |  |
| Noisy .3 | × | × | × | × |
| Noisy .4 | × | × | × | × |
| Noisy .5 | × | × | × | × |

#### Matrix entry similarity (F-score)

The binary F-score analysis (Figure 2) revealed that noisy data maintained perfect entry-wise agreement (100%) with the original datasets, as only a subset of values were replaced and the matrix sparsity was preserved. SE-generated synthetic datasets showed notably lower F-scores, particularly for `amgut2` (3.6%) and `MOMS-PI` (23%). In contrast, SP-generated synthetic datasets achieved relatively higher F-scores, comparable to those observed in the bootstrap datasets (e.g., 72% for `amgut1`, 68% for `amgut2`). These results are summarized in Table 3.

**Table 3:** Percentage of matching presence-absence entries between generated datasets and corresponding real microbiome datasets, reflecting data fidelity.

|  | amgut1 | amgut2 | GUT | MOMS-PI |
| --- | --- | --- | --- | --- |
| Bootstrap | 72 | 68 | 58 | 47 |
| Synthetic SE | 71 | 3.6 | 52 | 23 |
| Synthetic SP | 72 | 68 | 58 | 47 |
| Noisy | 100 | 100 | 100 | 100 |

**Figure 2:**
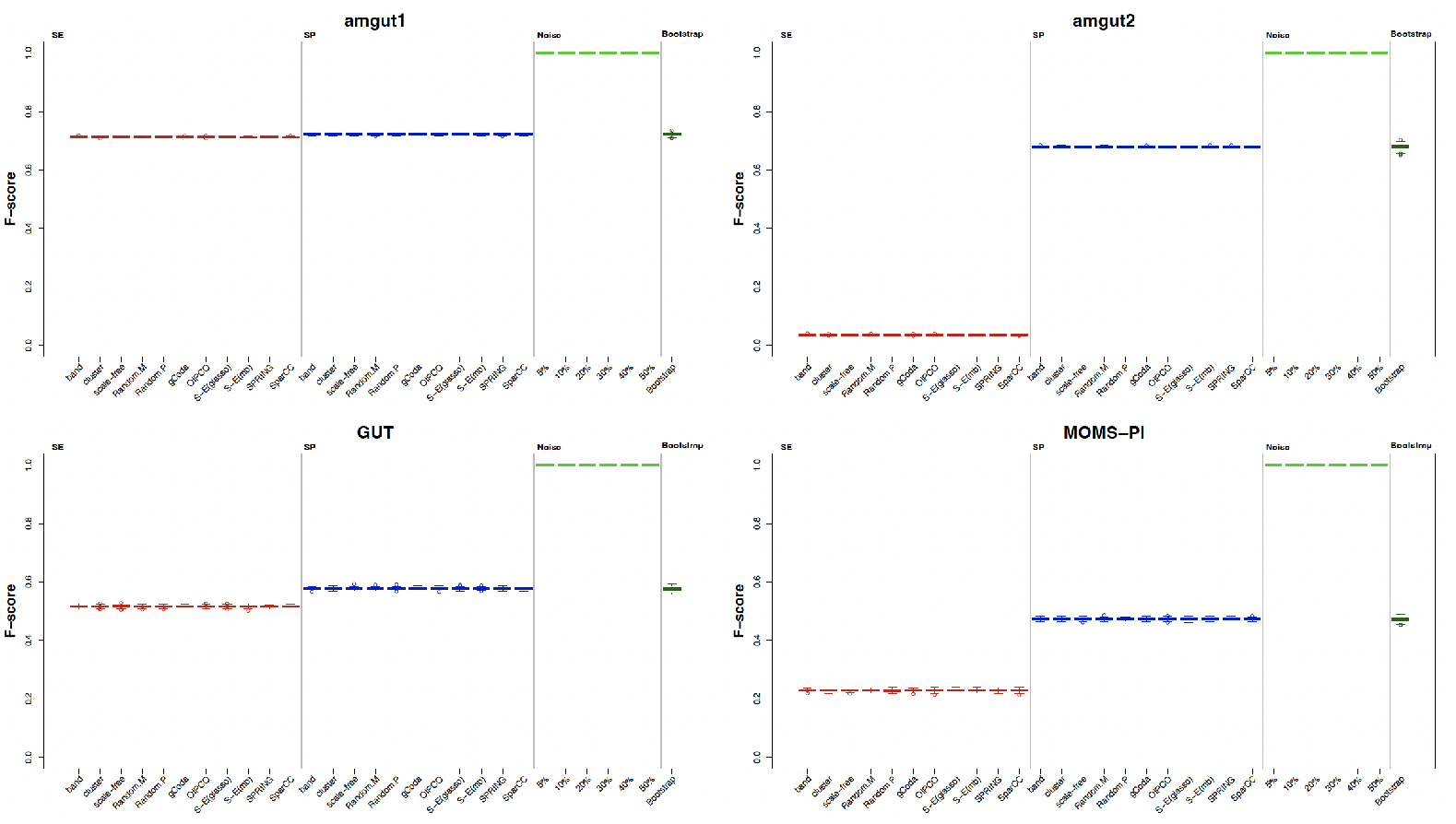
Box plots comparing the F-score for presence-absence agreement between real data and various generated datasets across four microbiome cohorts.

#### Distributional similarity (KS test)

As shown in Table 4, synthetic datasets generated by the SP method preserved OTU-level distributions exceptionally well, with none of the OTUs showing significant distributional differences across all datasets. Bootstrap datasets and noisy datasets with up to 10% perturbation also demonstrated complete distributional alignment. In contrast, SE-generated datasets showed substantial deviations: for example, 92% of OTUs in `amgut1` and 100% in `amgut2` were significantly different from the real data. Noisy datasets showed increasing divergence as noise levels rose, with 50% noise yielding 100% OTU-level differences in some cases. These results collectively indicate that the SE method produces synthetic data that deviates significantly from real data across all criteria, while the SP method, bootstrap samples, and noisy datasets (up to 10%) more closely preserve the statistical properties of the original microbiome data. For further details, refer to Figures S13 through S16.

**Table 4:**
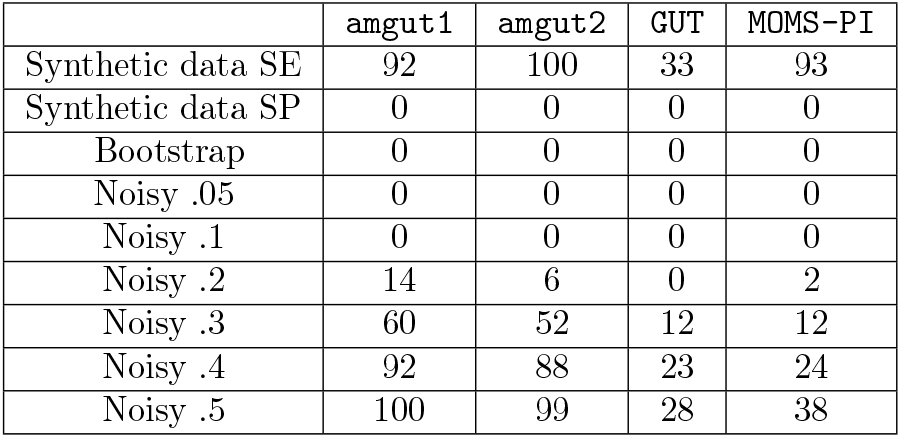
Percentage of OTUs exhibiting statistically significant differences in abundance distributions between generated and real datasets, based on Kolmogorov–Smirnov tests.

|  | amgut1 | amgut2 | GUT | MOMS-PI |
| --- | --- | --- | --- | --- |
| Synthetic data SE | 92 | 100 | 33 | 93 |
| Synthetic data SP | 0 | 0 | 0 | 0 |
| Bootstrap | 0 | 0 | 0 | 0 |
| Noisy .05 | 0 | 0 | 0 | 0 |
| Noisy .1 | 0 | 0 | 0 | 0 |
| Noisy .2 | 14 | 6 | 0 | 2 |
| Noisy .3 | 60 | 52 | 12 | 12 |
| Noisy .4 | 92 | 88 | 23 | 24 |
| Noisy .5 | 100 | 99 | 28 | 38 |

### 3.2 Algorithms performance on constructing microbiome networks based on generated data

We evaluated the performance of six microbial network inference algorithms (gCoda, OIPCQ, S-E(glasso), S-E(mb), SPRING, and SparCC) across four real microbiome datasets (`amgut1`, `amgut2`, `GUT`, and `MOMS-PI`). We generated three types of datasets derived from the real data: synthetic data, noisy data, and bootstrap data (described in Section 2.2). In each case, networks inferred from generated data were compared to appropriate reference networks, using F-score as the primary metric (See section 2.4).

#### gCoda Results

Across all datasets, the F-scores of gCoda generally follow the trend: SE *<* SP *<* (noisy & bootstrap) (in terms of mean values). For `amgut1`, `amgut2`, and `MOMS-PI`, noise levels of 5%, 10%, and 20% yield the highest F-scores. For `GUT`, only 5% and 10% noise levels perform best. Notably, the results of gCoda show a lack of significant difference across different synthetic topologies (e.g., band, cluster, and random) for certain datasets, suggesting limited sensitivity of the algorithm to underlying network structures. In `amgut1` the Random.M topology and the gCoda-inferred topology from SP are not significantly different. In `amgut2`, the band topology generated by SE and the gCoda-inferred topology from SE are not significantly different, implying convergence to a used training topology. In `GUT`, the gCoda-based topology from SP data shows no significant difference with noise levels of 40% and 50% (see Figure S28). These results highlight a potential issue: gCoda often yields statistically indistinguishable outcomes across very different data structures, even when real differences exist, questioning its discriminative power.

#### OIPCQ Results

For OIPCQ SE-generated data generally result in lower F-scores than SP-generated data, with synthetic data performing worse than both noisy and bootstrap datasets. Bootstrap results for `amgut1`, `amgut2`, and `MOMS-PI` are lower than 5%, 10%, and 20% noise levels; for `GUT`, they are only higher than 50% noise. In `amgut1`, the results of band topology from SP do not show a statistically significant difference at a 50% noise level. For `amgut2`, none of the synthetic or noisy datasets show statistically significant differences. In `GUT`, the bootstrap results even lower than 40% noise level. Also, the cluster topology derived from SE does not show a statistically significant difference compared to the OIPCQ-inferred topology from SE and the band and Random.M topologies from SP. Similar indistinguishability is observed in `MOMS-PI` between SE band and SP Random.P topologies (see Figure S29). These findings suggest that while OIPCQ can distinguish noise from clean data, it may not fully differentiate between subtle topological variations in synthetic data.

#### S-E(glasso) Results

For `amgut1`, `amgut2`, and `MOMS-PI`, SE-based synthetic data yields lower F-scores than SP-based data. For `GUT`, SE slightly outperforms SP. For `amgut2`, SE results approach zero due to only 3.6% entry overlap with the real data. Noise levels of 5% and 10% show the highest F-scores for most datasets, with all noise levels outperforming bootstrap in `GUT`. In `amgut1`, synthetic SE band data is not significantly different from SP Random.P data or noise levels of 30% and 40%. Similarly, multiple topologies yield statistically indistinguishable results in `amgut2`, `GUT`, and `MOMS-PI`, including random and algorithm-derived topologies (see Figure S30). These results indicate that S-E(glasso) is moderately sensitive to synthetic data quality.

#### S-E(mb) Results

S-E(mb) exhibits a diverse performance range. For `amgut1`, some synthetic datasets outperform noisy data. For `GUT`, noisy data consistently outperforms both synthetic and bootstrap datasets. Multiple topologies yield non-significant results in `amgut1`, `amgut2`, `GUT`, and `MOMS-PI`, suggesting limited sensitivity of the algorithm to differences in network structure (see Figure S31). Bootstrap results in `MOMS-PI` are not significantly different from 20% noise. These observations raise concerns about S-E(mb)’s ability to reliably distinguish between networks with meaningful topological structure (e.g., clustered or inferred from data) and random-like graphs.

#### SPRING Results

SPRING results vary considerably by dataset. For `GUT`, most results from SE and SP synthetic datasets show no statistically significant differences. For `amgut1` SE-based synthetic data outperform SP-based data; the reverse holds for `amgut2` and `MOMS-PI`. In some cases (e.g., `amgut1`, `amgut2`), synthetic data outperforms noisy and bootstrap data, a surprising outcome. Many topologies yield non-significant differences in `amgut1`, `amgut2`, `GUT`, and `MOMS-PI`. For example, in `MOMS-PI`, cluster and Random.P topologies from SP show indistinguishable results, potentially indicating robustness or insensitivity to structure (see Figure S32). For `MOMS-PI` and `GUT`, the results of all noisy datasets are higher than the bootstrap and for `GUT`, bootstrap results are below 0.5.

#### SparCC Results

SparCC shows strong performance on noisy and bootstrap data across all datasets. For `amgut2` and `MOMS-PI`, SE-based synthetic data yield NaN values for almost all datasets (SparCC returns NaN when it deems the inferred correlation results to be statistically unreliable.) SP-based data generally perform worse than noisy and bootstrap data. For `amgut1` and `GUT`, noise levels of 5% and 10% outperform bootstrap. Several synthetic topologies produce statistically indistinguishable results in `amgut1`, `amgut2`, `GUT`, and `MOMS-PI`, including random and algorithm-derived networks, further questioning the utility of topology-driven synthetic evaluations (see Figure S33). Figure 3 shows the violin plots of the F-score for each algorithm on `amgut1`. Refer to Figures S17 to S27 for additional violin plots of Precision, Recall, and F-score across all real datasets.

**Figure 3:**
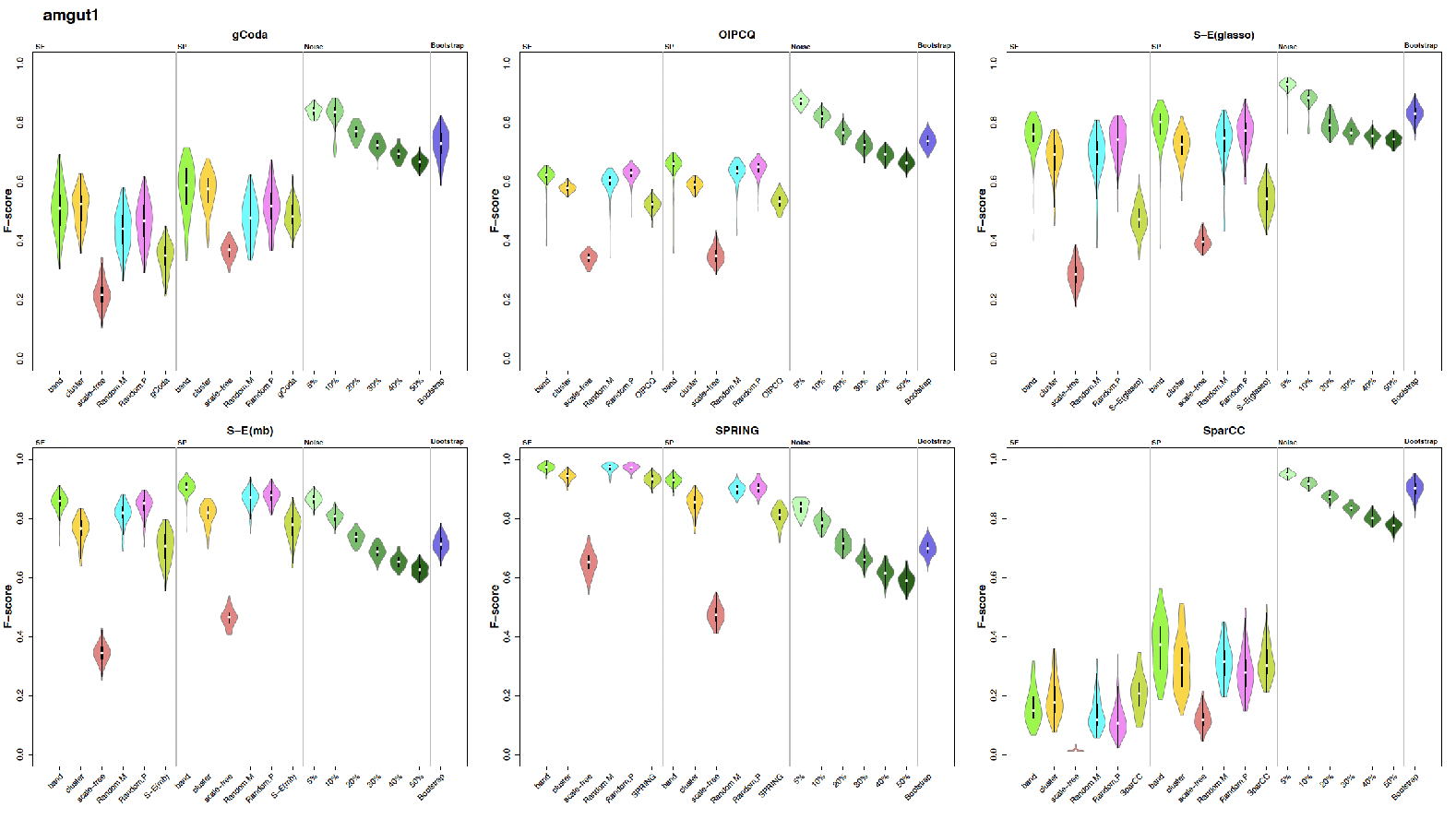
Violin plots showing the distribution of F-scores for six network inference algorithms applied to the amgut1 dataset across synthetic (SE and SP), noisy, and bootstrap data.

### 3.3 Comparing the results of all algorithms across all generated data from real-world datasets

#### amgut1

For this real dataset, the best results for all synthetic datasets are obtained from the SPRING algorithm. However, for noisy and bootstrap data, the best results are obtained from the SparCC algorithm (see Figure S34).

#### amgut2

For this real dataset, the SPRING algorithm yields the best results for all synthetic SP-generated data from specified topologies. For SP-generated synthetic data from algorithm topologies, SPRING provides the best results but does not show statistically significant differences compared to the S-E(mb) method. For SE-generated synthetic data, the results are very low, and often there is no statistically significant difference between the highest results (the results of the gCoda algorithm do not show statistically significant differences compared to the OIPCQ and SPRING algorithms). For SE-generated synthetic data from algorithm topologies, gCoda provides the best results. For 5% noise and bootstrapping, SparCC gives the best result. For 10% noise, SparCC results are the best and do not show statistically significant differences compared to gCoda and OIPCQ. For 20% to 50% noise, the OIPCQ algorithm yields the best result (see Figure S35).

#### GUT

For this real dataset, the OIPCQ algorithm provides the best results for all synthetic data with specified topologies. For synthetic data resulting from algorithm topologies, gCoda yields the best result. For noise levels of 5% and 10%, S-E(mb) performs best, and for other noise levels, the SPRING algorithm has the best performance (see Figure S36).

#### MOMS-PI

For this real dataset, among all SP-based synthetic data from the given topologies, the best result comes from the SPRING algorithm. For SE-based synthetic data from the given topologies, the best solution is provided by the OIPCQ algorithm. For both SE and SP synthetic data from the topologies of the algorithms, the best result is achieved by the gCoda algorithm. For data with 5% noise, the best result is obtained from the S-E(glasso) algorithm. For other noise levels, SparCC performs best. For bootstrap data, the highest results come from the S-E(glasso) and SparCC algorithms, which are not statistically significantly different (see Figure 4).

**Figure 4:**
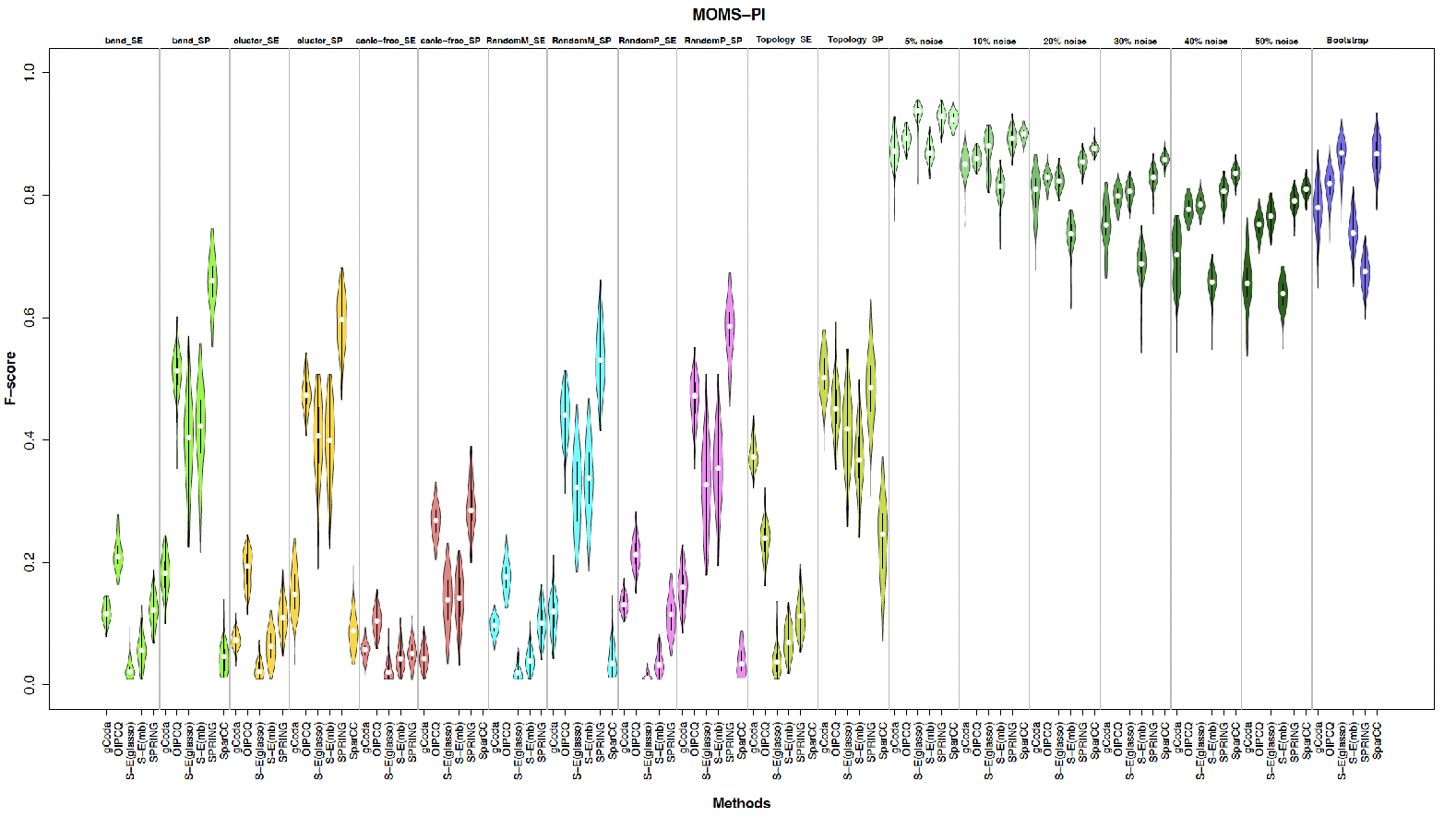
Violin plots comparing F-scores of each microbial network inference algorithm across generated datasets from the `MOMS-PI` cohort. *“band SE”* refers to datasets generated using the band topology with the SE method. Similarly, *“Topology SE”* denotes datasets generated using each algorithm’s inferred topology with the SE method. These comparisons highlight algorithm robustness and performance consistency across diverse data generation strategies.

### 3.4 Summary observations across all algorithms

Our comprehensive evaluation across all datasets and data generation strategies reveals several consistent and critical trends in algorithm behavior. Notably, synthetic data generated from scale-free topologies consistently produced the lowest F-scores across all methods, highlighting the limitations of using this commonly assumed structure in benchmarking microbial networks. Despite being frequently used, scale-free topologies appear to poorly reflect the complexity of real microbial ecosystems, leading to artificially depressed performance metrics. Among all algorithms, SparCC exhibited the most robust and reproducible performance on bootstrap datasets, with only a marginal and statistically insignificant advantage by S-E(glasso) in the `MOMS-PI` dataset. This reinforces SparCC’s resilience to sampling variability and supports its use in reproducibility-focused assessments. Interestingly, many algorithms failed to significantly differentiate between structured and unstructured synthetic topologies—including banded, clustered, random, and even real-data-inferred networks. This structural insensitivity undermines confidence in evaluations based solely on synthetic benchmarks and calls for stronger criteria rooted in data realism and biological plausibility. Despite clear statistical distinctions between datasets generated via the SE and SP methods (with SE often diverging substantially from real data in abundance distribution and diversity), some algorithms did not exhibit statistically significant performance differences when applied to SE versus SP datasets. This further questions their ability to respond appropriately to changes in data structure and statistical properties. Moreover, and perhaps most strikingly, both bootstrap and low-noise data sets (10%) consistently yielded higher F-scores than synthetic datasets, even when the latter were derived from real-data-inferred topologies. This was especially pronounced in the `GUT` and `MOMS-PI` datasets, where the F-score distributions for noisy datasets (processed by S-E(mb) and SPRING) were substantially higher than those for bootstrap data, with no overlapping ranges. Given that bootstrap data are inherently closer to the original datasets, this finding is counterintuitive and emphasizes the need for caution when interpreting high synthetic performance as a proxy for robustness. These observations collectively underscore the importance of reproducibility-based evaluation using real-data-derived perturbations, and they advocate for greater scrutiny of traditional synthetic benchmarks in microbiome network inference. Figure 5 presents the F-score distribution across all algorithms on bootstrap datasets, offering a comparative view of algorithmic reproducibility under realistic sampling variability.

**Figure 5:**
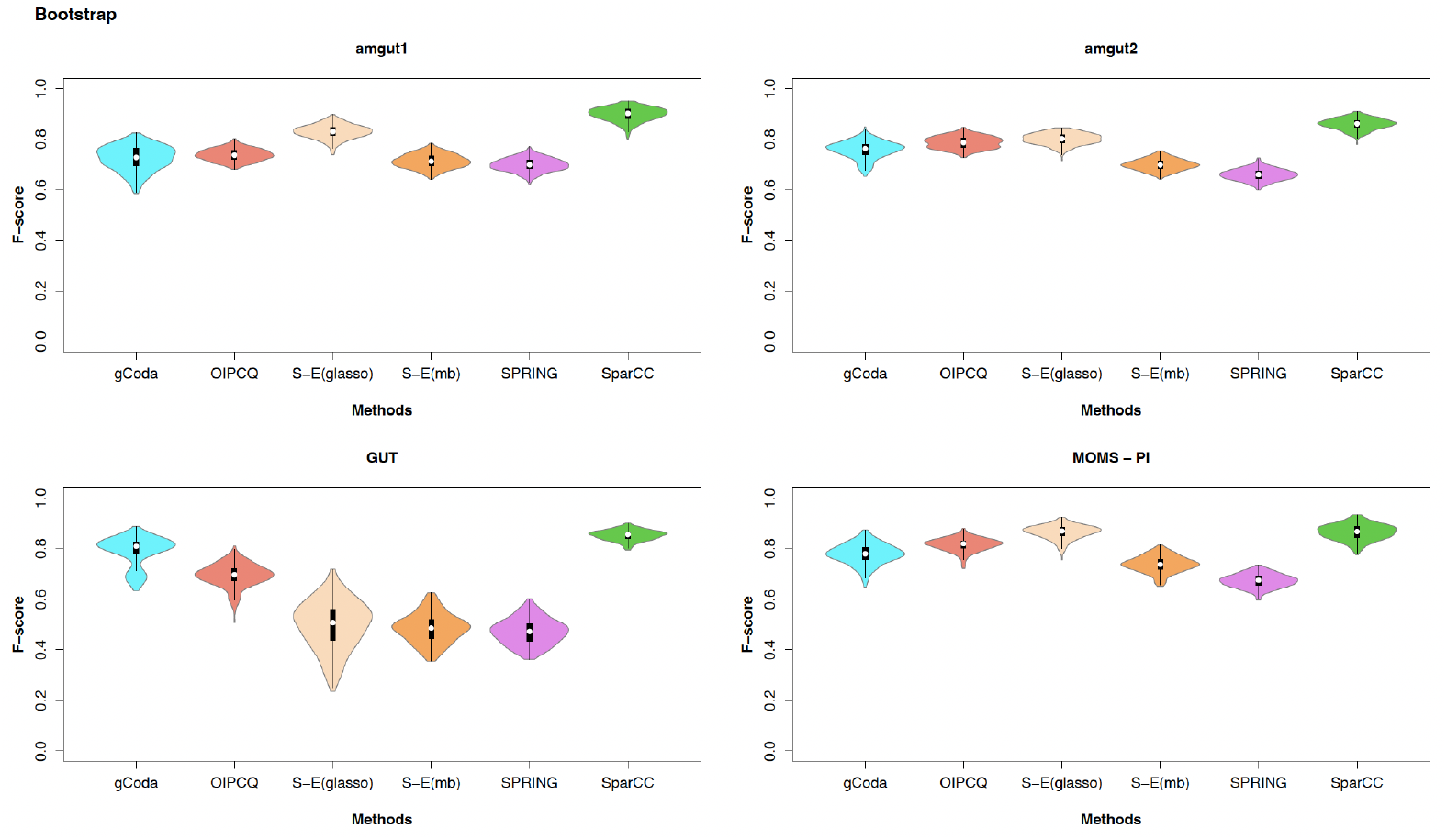
Violin plots comparing the reproducibility of six microbial network inference algorithms on bootstrap datasets across all four real microbiome datasets.

## 4 Discussion

In this study, we present a comprehensive and reproducibility-centered evaluation framework for microbial network inference, designed to address a central challenge in microbiome research: the lack of an experimentally validated ground truth. By benchmarking six widely used algorithms across both real and variously perturbed datasets, we demonstrate that reproducibility, measured through performance stability on bootstrap and low-noise variants of real data, can serve as a practical and rigorous proxy for validation. Our findings reveal that while synthetic datasets remain a common benchmarking tool, they often fail to preserve the key statistical and structural properties of real microbiome data. This limitation is especially pronounced for synthetic data generated using the SE method, which frequently deviates in OTU distributions, matrix sparsity, and diversity indices. Although synthetic networks derived from real-data-inferred topologies might appear more biologically grounded, our results show that even these often yield inflated or misleading performance metrics. Notably, in several instances, synthetic datasets generated from random or biologically implausible topologies yielded higher F-scores than those derived from realistic perturbations of real data—an outcome that calls into question the reliability of synthetic benchmarking when applied without careful validation. In contrast, bootstrap and mildly noisy datasets (≤ 10% perturbation) preserve the compositional and structural characteristics of real data and expose algorithmic weaknesses that synthetic datasets fail to reveal. This highlights a key insight: reproducibility under real-world variability offers a more faithful test of algorithm robustness than performance on idealized simulations. Algorithms that perform consistently on bootstrap and noisy data are more likely to generalize across biological contexts and sample fluctuations. Our evaluation also exposes a common shortcoming among many algorithms: structural insensitivity. Several methods failed to meaningfully distinguish between realistic and randomly structured synthetic networks. This raises concerns about the ecological interpretability of inferred networks and suggests that performance metrics such as F-score, when computed solely on synthetic benchmarks, may not reliably reflect biological plausibility. We therefore advocate a shift in benchmarking practices for microbial network inference. When synthetic data are used, their statistical fidelity to the original dataset must be explicitly validated using quantitative tests (e.g., KS test, matrix entry similarity, diversity comparisons). More importantly, we propose that reproducibility on data derived from real-world samples via bootstrapping and controlled noise should be prioritized as a core evaluation strategy. Our proposed pipeline provides a step-by-step methodology for doing so, grounded in both theoretical rigor and empirical insight.

Beyond microbiome studies, these results have implications for other domains in computational biology where ground truth is unavailable or uncertain. As the field moves toward more complex and integrated microbial ecosystem modeling, reproducibility will not only enhance method selection but also contribute to the reliability of downstream biological inferences. In summary, we argue that reproducibility is not a secondary metric, but a necessary criterion for evaluating microbial network inference algorithms. Our findings challenge the overreliance on synthetic topologies and advocate for biologically and statistically grounded benchmarks that reflect the realities of microbial data. By adopting reproducibility as a central pillar of evaluation, the field can develop and validate methods that are both technically sound and ecologically meaningful.

## 5 Conclusion

This study addresses a fundamental challenge in microbial network inference: how to reliably evaluate algorithm performance in the absence of a true gold standard. By benchmarking six widely used methods on real microbiome data and a range of biologically grounded perturbations—including bootstrap resampling and controlled noise—we demonstrate that traditional synthetic benchmarks frequently fail to capture the statistical and ecological complexity of real data. Our findings reveal that reproducibility under realistic data variation is a more meaningful and reliable evaluation strategy than performance on idealized synthetic datasets. Algorithms that perform consistently on perturbed versions of real data are more likely to generalize across experimental contexts, offering stronger ecological validity. To guide future evaluations, we propose a reproducibility-centered workflow for benchmarking new microbial network inference methods—designed to assess algorithmic robustness by emphasizing real-data perturbations and requiring statistical validation of any synthetic benchmarks.

1. **Baseline Inference:** Apply the algorithm to a real microbiome dataset to construct a baseline interaction network.
2. **Bootstrap Generation:** Generate bootstrap datasets by resampling samples (rows) with replacement to introduce sampling variability while preserving ecological structure.
3. **Noisy Data Generation:** Introduce mild random noise (5–10%) to simulate technical or biological perturbations while maintaining compositional integrity. This level of noise simulates realistic variability without fundamentally altering the core patterns or balance within the dataset.
4. **Apply Algorithm to Perturbed Data:** Infer networks from each bootstrap and noisy replicate using the same algorithm.
5. **Evaluate Across Data Types:** Compare algorithm performance across bootstrap, noisy, and synthetic datasets. When using synthetic data, generate it using the algorithm’s own inferred topology and validate its realism via diversity metrics, matrix entry similarity, and OTU-wise distributional alignment (e.g., KS test). This allows for nuanced interpretation of how algorithmic robustness degrades as data diverge from biological reality.

This framework offers a statistically rigorous, reproducibility-focused alternative to synthetic-only evaluations. It empowers researchers to assess methodological reliability under real-world variability and supports the development of more robust and biologically interpretable network inference tools.

While synthetic data benchmarks remain valuable, they should be used cautiously and validated thoroughly—such as by comparing synthetic datasets’ ecological properties to real data—so that their limitations are recognized. By adopting reproducibility as a core benchmarking principle, we can advance the field toward more trustworthy and ecologically informed microbiome network analyses.

## Data availability

The datasets and codes can be found in the GitHub repository https://github.com/zghaeli/Microbiome-Network-Evaluation

## Author contributions

Z.G. conducted the core research and data analysis. R.A. and C.E. provided guidance on study design, methodology, and interpretation of results. Both R.A. and C.E. contributed to the conceptual framework and assisted with manuscript preparation. All authors contributed to drafting, revising, and approving the final manuscript.

## Additional information

**Supplementary Information** The online version contains supplementary material available at

**Correspondence** and requests for materials should be addressed to C.E.

## References

1. Berg, G. et al. Microbiome definition re-visited: old concepts and new challenges. Microbiome 8, 1–22 (2020).

2. Aggarwal, N. et al. Microbiome and human health: current understanding, engineering, and enabling technologies. Chemical reviews 123, 31–72 (2022).

3. Cox, M. J., Cookson, W. O. & Moffatt, M. F. Sequencing the human microbiome in health and disease. Human molecular genetics 22, R88–R94 (2013).

4. Structure, function and diversity of the healthy human microbiome. nature 486, 207–214 (2012).

5. Xia, C. et al. Human microbiomes in cancer development and therapy. MedComm 4, e221 (2023).

6. Qasem, H. H. & El-Sayed, W. M. The bacterial microbiome and cancer: development, diagnosis, treatment, and future directions. Clinical and Experimental Medicine 25, 12 (2024).

7. Martinelli, F. et al. Whole-body metabolic modelling reveals microbiome and genomic interactions on reduced urine formate levels in Alzheimer’s disease. Scientific Reports 14, 6095 (2024).

8. Zhou, K., Baranova, A., Cao, H., Sun, J. & Zhang, F. Gut microbiome and schizophrenia: insights from two-sample Mendelian randomization. Schizophrenia 10, 75 (2024).

9. Faust, K. & Raes, J. Microbial interactions: from networks to models. Nature Reviews Microbiology 10, 538–550 (2012).

10. Abegaz, F. et al. A strategy for differential abundance analysis of sparse microbiome data with group-wise structured zeros. Scientific Reports 14, 12433 (2024).

11. Silverman, J. D., Roche, K., Mukherjee, S. & David, L. A. Naught all zeros in sequence count data are the same. Computational and structural biotechnology journal 18, 2789–2798 (2020).

12. Aitchison, J. A new approach to null correlations of proportions. Journal of the International Association for Mathematical Geology 13, 175–189 (1981).

13. Kurtz, Z. D. et al. Sparse and compositionally robust inference of microbial ecological networks. PLoS computational biology 11, e1004226 (2015).

14. Gorstein, E., Aghdam, R. & Solís-Lemus, C. HighDimMixedModels. jl: Robust high-dimensional mixed-effects models across omics data. PLOS Computational Biology 21, e1012143 (2025).

15. Nelson, R., Aghdam, R. & Solis-Lemus, C. MiNAA: Microbiome Network Alignment Algorithm. Journal of Open Source Software 9, 5448 (2024).

16. Berry, D. & Widder, S. Deciphering microbial interactions and detecting keystone species with co-occurrence networks. Frontiers in microbiology 5, 219 (2014).

17. Banerjee, S., Schlaeppi, K. & Van Der Heijden, M. G. Keystone taxa as drivers of microbiome structure and functioning. Nature Reviews Microbiology 16, 567–576 (2018).

18. Friedman, J. & Alm, E. J. Inferring Correlation Networks from Genomic Survey Data. PLoS Comput Biol 8, e1002687. 10.1371/journal.pcbi.1002687 (2012).

19. Fang, H., Huang, C., Zhao, H. & Deng, M. CCLasso: correlation inference for compositional data through Lasso. Bioinformatics 31, 3172–3180 (2015).

20. Ban, Y., An, L. & Jiang, H. Investigating microbial co-occurrence patterns based on metagenomic compositional data. Bioinformatics 31, 3322–3329 (2015).

21. Yoon, G., Gaynanova, I. & Müller, C. L. Microbial networks in SPRING-Semi-parametric rank-based correlation and partial correlation estimation for quantitative microbiome data. Frontiers in genetics 10, 516 (2019).

22. Faust, K. & Raes, J. CoNet app: inference of biological association networks using Cytoscape. F1000Research 5, 1519 (2016).

23. Yang, P., Yu, S., Cheng, L. & Ning, K. Meta-network: optimized species-species network analysis for microbial communities. BMC genomics 20, 143–151 (2019).

24. Jiang, S. et al. HARMONIES: a hybrid approach for microbiome networks inference via exploiting sparsity. Frontiers in Genetics 11, 445 (2020).

25. Tackmann, J., Rodrigues, J. F. M. & von Mering, C. Rapid inference of direct interactions in large-scale ecological networks from heterogeneous microbial sequencing data. Cell systems 9, 286–296 (2019).

26. Feng, C. et al. MicroNet-MIMRF: a microbial network inference approach based on mutual information and Markov random fields. Bioinformatics Advances 4, vbae167 (2024).

27. Fang, H., Huang, C., Zhao, H. & Deng, M. gCoda: conditional dependence network inference for compositional data. Journal of Computational Biology 24, 699–708 (2017).

28. Aghdam, R., Shan, S., Lankau, R. & Solis-Lemus, C. Leveraging Bayesian Networks for Consensus Network Construction and Multi-Method Feature Selection to Decode Disease Prediction. bioRxiv, 2025–04 (2025).

29. Yooseph, S. et al. Stool microbiota composition is associated with the prospective risk of Plasmodium falciparum infection. BMC genomics 16, 1–15 (2015).

30. Tavakoli, S. & Yooseph, S. Learning a mixture of microbial networks using minorization–maximization. Bioinformatics 35, i23–i30 (2019).

31. Osborne, N., Peterson, C. B. & Vannucci, M. Latent network estimation and variable selection for compositional data via variational EM. Journal of Computational and Graphical Statistics 31, 163–175 (2022).

32. Cario, M. C. & Nelson, B. L. Modeling and generating random vectors with arbitrary marginal distributions and correlation matrix tech. rep. (Technical Report, Department of Industrial Engineering and Management …, 1997).

33. Csardi, G. & Nepusz, T. The igraph software. Complex syst 1695, 1–9 (2006).

34. Cassol, I., Ibañez, M. & Bustamante, J. P. Key features and guidelines for the application of microbial alpha diversity metrics. Scientific Reports 15, 622 (2025).

35. Dixon, P. VEGAN, a package of R functions for community ecology. Journal of vegetation science 14, 927–930 (2003).

