## Supplementary Materials for "Evaluating microbial network inference methods: Moving beyond synthetic data with reproducibility-driven benchmarks"

#### 1 Comparison of Diversity Indices Between Real-World and Generated (Synthetic, Noisy, Bootstrap) Datasets

##### 1.1 Shannon Index

For each sample of the real dataset, we calculate the Shannon index, resulting in a vector sized according to the number of real data samples. Similarly, we compute the Shannon index for each generated dataset. To identify significant differences between each dataset and the real data, we utilize the p-value from the Wilcoxon test. Given that we have 100 repetitions for each type of noisy and synthetic data, we create a violin plot of these 100 p-values. We also generate this plot for bootstrap data, which matches the sample size of the real data. Figure S1 illustrates the p-values for all datasets derived from the `amgut1` real data.

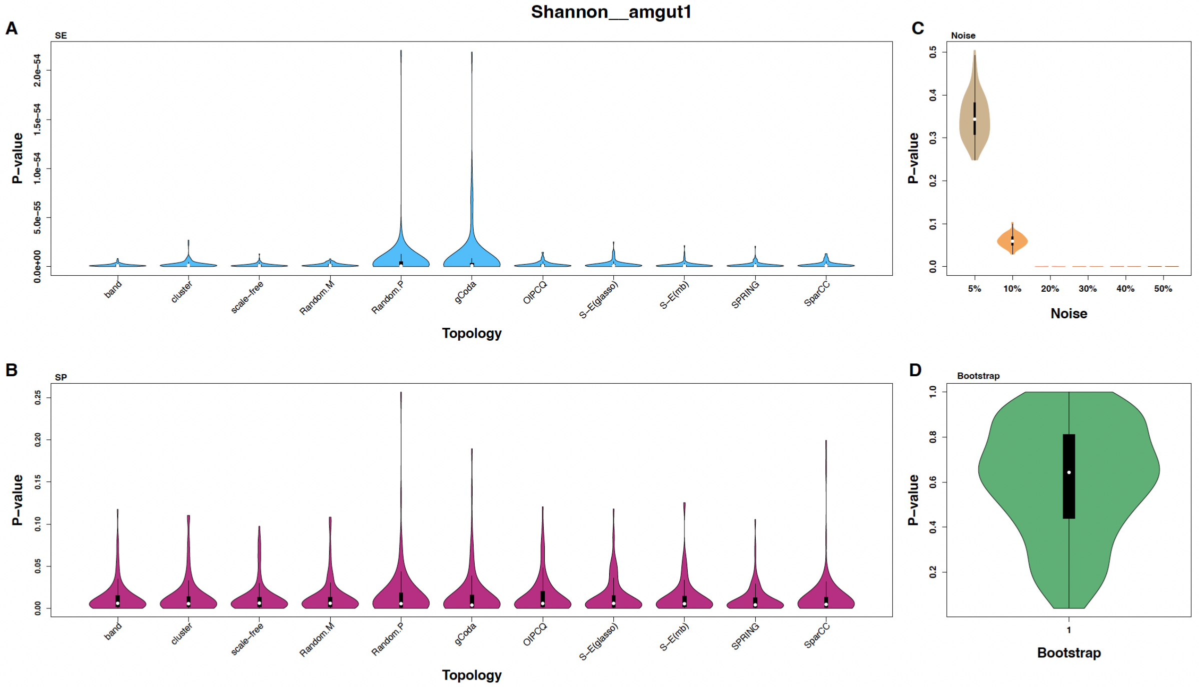


Figure S1: Violin plots showing the distribution of p-values from Wilcoxon tests comparing the Shannon index between the `amgut1` dataset and multiple generated datasets. Panels display results for (A) SE-generated synthetic datasets, (B) SP-generated synthetic datasets, (C) noisy datasets with varying noise levels, and (D) bootstrap datasets.

The p-values for all synthetic datasets are below 0.05, indicating that these data significantly differ from the `amgut1` data regarding the Shannon index. Furthermore, the synthetic data from the SE method demonstrate a greater difference compared to the SP data. Additionally, the datasets with 20 %, 30 %, 40 %, and 50 % noise also show significant differences, although the magnitude of their differences is less than that of the SE data. In contrast, the data with 5 % and 10 % noise, along with the bootstrap data, do not exhibit significant differences according to the Wilcoxon test.

The results of the Wilcoxon test for Shannon values in the `amgut2`, `GUT`, and `MOMS-PI` datasets are shown in Figures S2, S3, and S4, respectively.

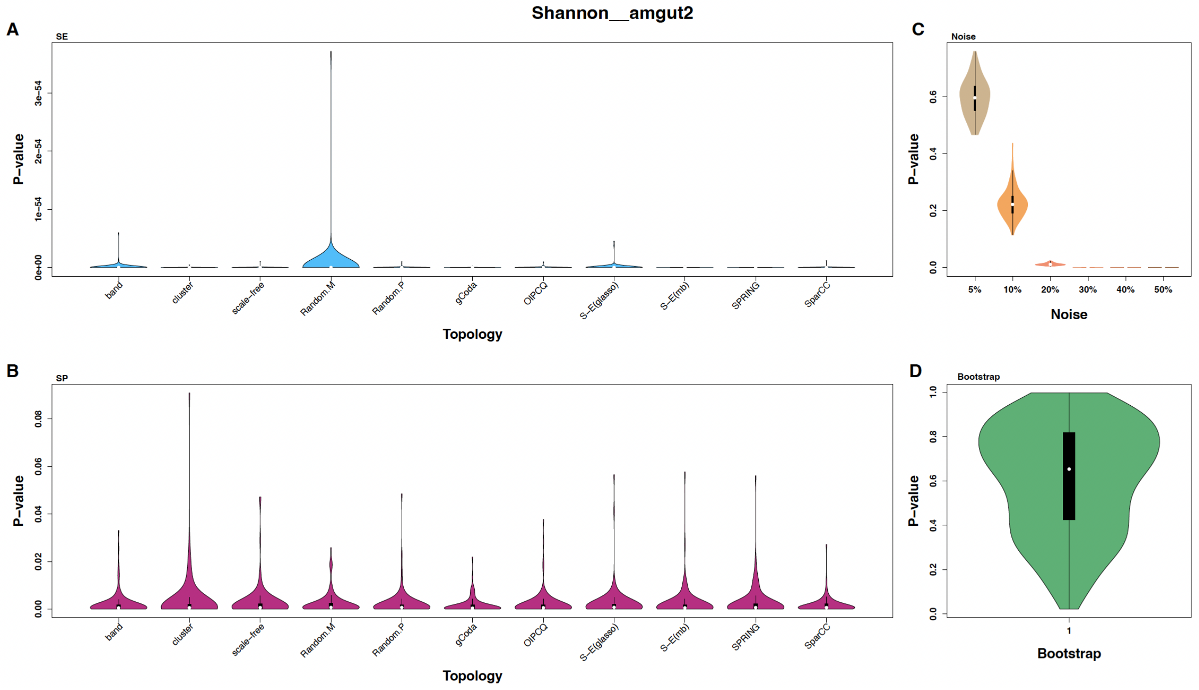


Figure S2: Violin plots showing the distribution of p-values from Wilcoxon tests comparing the Shannon index between the `amgut2` dataset and multiple generated datasets. Panels display results for (A) SE-generated synthetic datasets, (B) SP-generated synthetic datasets, (C) noisy datasets with varying noise levels, and (D) bootstrap datasets.

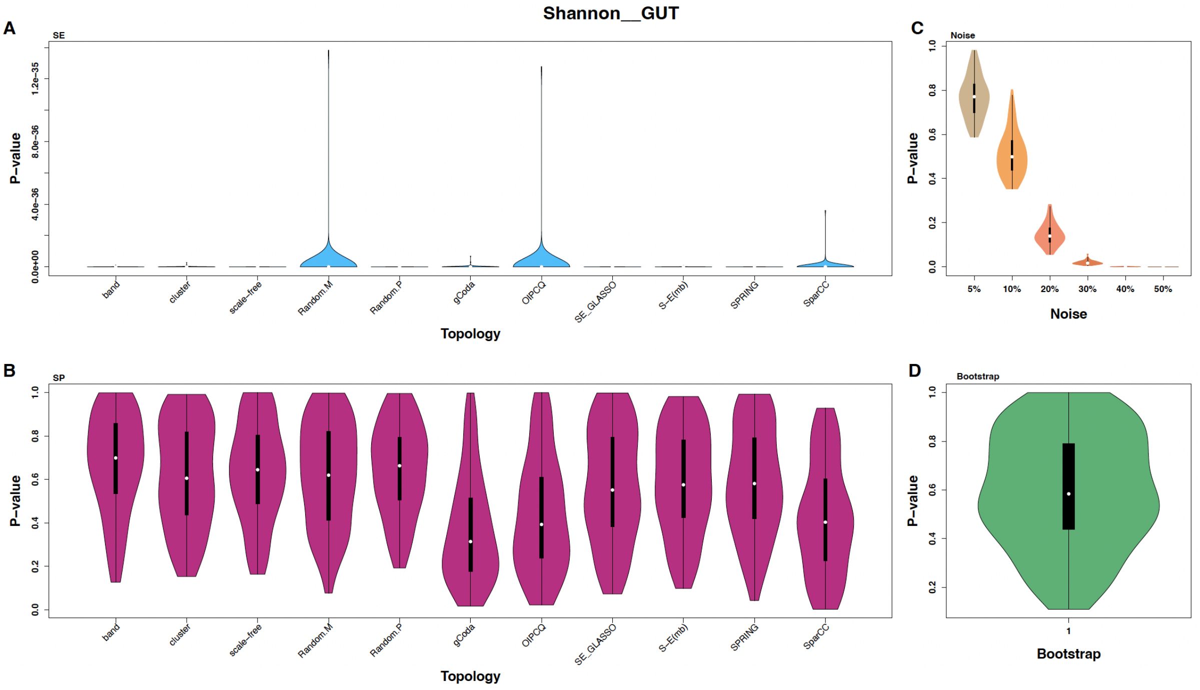


Figure S3: Violin plots showing the distribution of p-values from Wilcoxon tests comparing the Shannon index between the `GUT` dataset and multiple generated datasets. Panels display results for (A) SE-generated synthetic datasets, (B) SP-generated synthetic datasets, (C) noisy datasets with varying noise levels, and (D) bootstrap datasets.

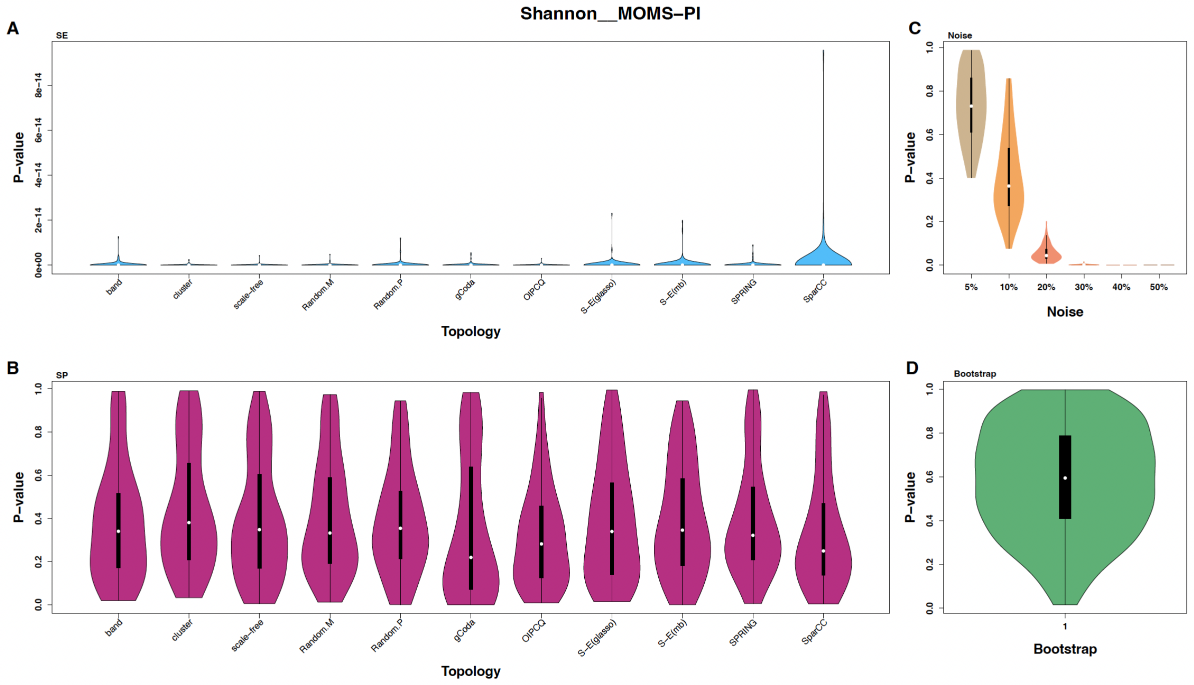


Figure S4: Violin plots showing the distribution of p-values from Wilcoxon tests comparing the Shannon index between the MOMS-PI dataset and multiple generated datasets. Panels display results for (A) SE-generated synthetic datasets, (B) SP-generated synthetic datasets, (C) noisy datasets with varying noise levels, and (D) bootstrap datasets.

#### 1.2 InvSimpson Index

For each sample of the real dataset, we calculate the InvSimpson index, resulting in a vector sized according to the number of real data samples. Similarly, we compute the InvSimpson index for each generated dataset. To identify significant differences between each dataset and the real data, we utilize the p-value from the Wilcoxon test. Given that we have 100 repetitions for each type of noisy and synthetic data, we create a violin plot of these 100 p-values. We also generate this plot for bootstrap data, which matches the sample size of the real data. Figure S5 illustrates the p-values for all datasets derived from the `amgut1` real data.

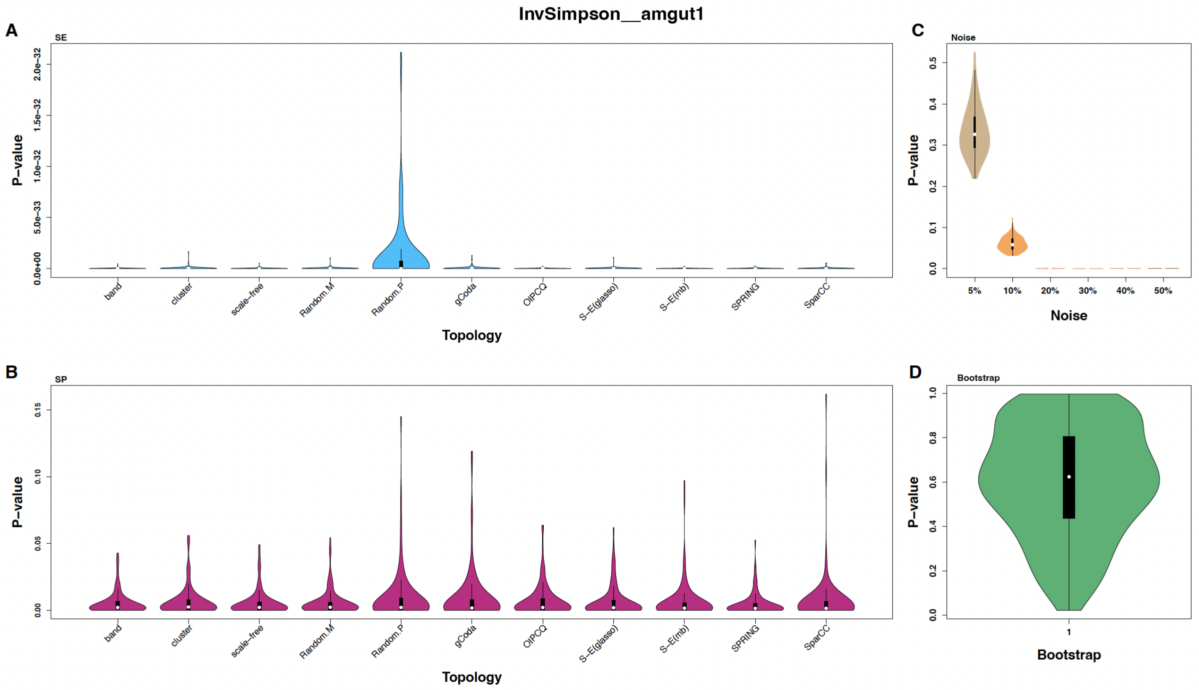


Figure S5: Violin plots showing the distribution of p-values from Wilcoxon tests comparing the InvSimpson index between the `amgut1` dataset and multiple generated datasets. Panels display results for (A) SE-generated synthetic datasets, (B) SP-generated synthetic datasets, (C) noisy datasets with varying noise levels, and (D) bootstrap datasets.

The p-values for all synthetic datasets are below 0.05, indicating that these data significantly differ from the `amgut1` data regarding the InvSimpson index. Furthermore, the synthetic data from the SE method demonstrate a greater difference compared to the SP data. Additionally, the datasets with 20 %, 30 %, 40 %, and 50 % noise also show significant differences, although the magnitude of their differences is less than that of the SE data. In contrast, the data with 5 % and 10 % noise, along with the bootstrap data, do not exhibit significant differences according to the Wilcoxon test.

The results of the Wilcoxon test for InvSimpson values in the `amgut2`, `GUT`, and `MOMS-PI` datasets are shown in Figures S6, S7, and S8, respectively.

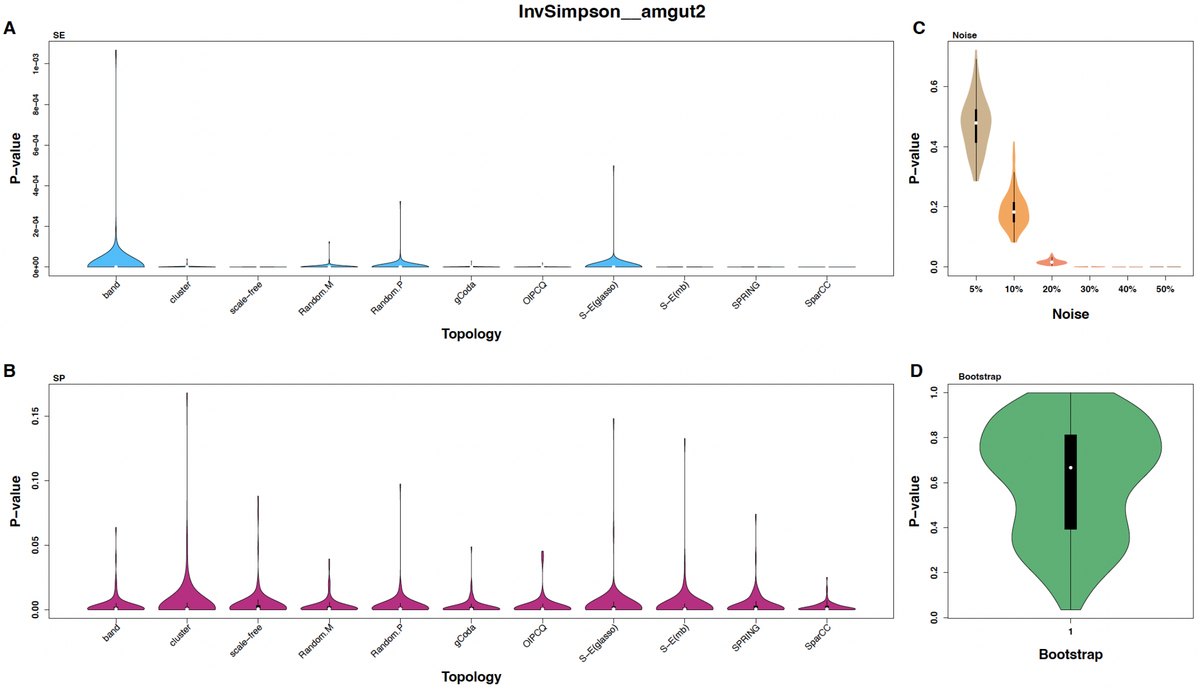


Figure S6: Violin plots showing the distribution of p-values from Wilcoxon tests comparing the InvSimpson index between the `amgut2` dataset and multiple generated datasets. Panels display results for (A) SE-generated synthetic datasets, (B) SP-generated synthetic datasets, (C) noisy datasets with varying noise levels, and (D) bootstrap datasets.

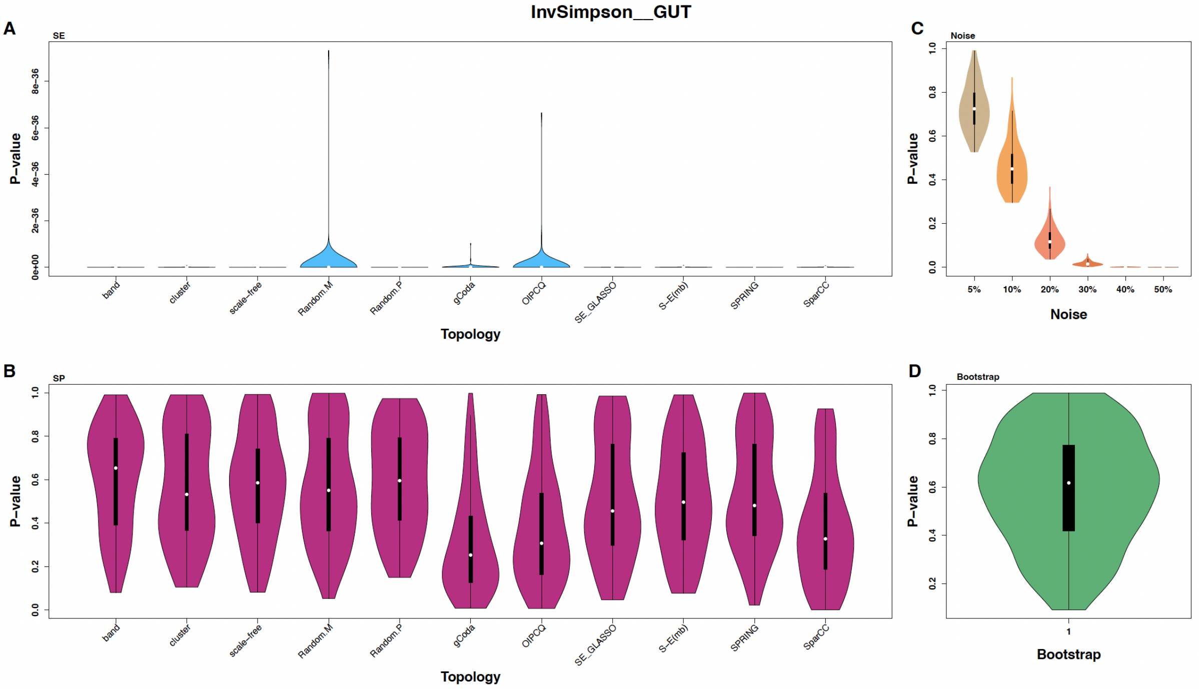


Figure S7: Violin plots showing the distribution of p-values from Wilcoxon tests comparing the InvSimpson index between the `GUT` dataset and multiple generated datasets. Panels display results for (A) SE-generated synthetic datasets, (B) SP-generated synthetic datasets, (C) noisy datasets with varying noise levels, and (D) bootstrap datasets.

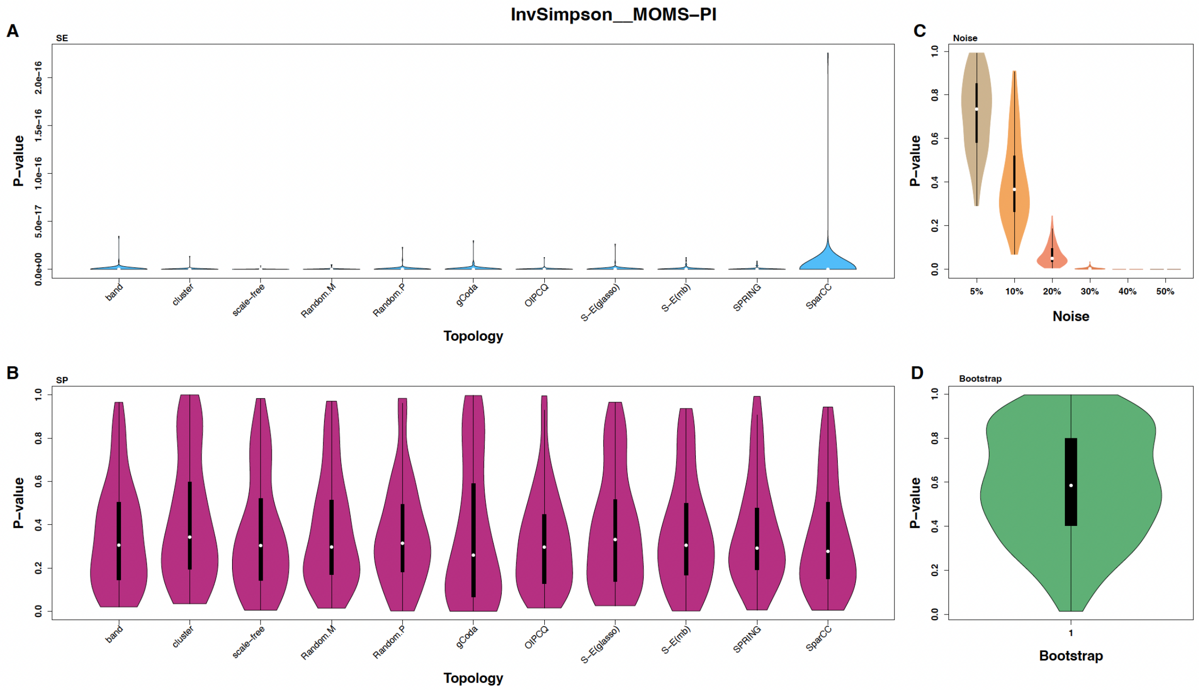


Figure S8: Violin plots showing the distribution of p-values from Wilcoxon tests comparing the InvSimpson index between the `MOMS-PI` dataset and multiple generated datasets. Panels display results for (A) SE-generated synthetic datasets, (B) SP-generated synthetic datasets, (C) noisy datasets with varying noise levels, and (D) bootstrap datasets.

#### 1.3 Richness Index

For each sample of the real dataset, we calculate the Richness index, resulting in a vector sized according to the number of real data samples. Similarly, we compute the Richness index for each generated dataset. To identify significant differences between each dataset and the real data, we utilize the p-value from the Wilcoxon test. Given that we have 100 repetitions for each type of noisy and synthetic data, we create a violin plot of these 100 p-values. We also generate this plot for bootstrap data, which matches the sample size of the real data. Figure S9 illustrates the p-values for all datasets derived from the `amgut1` real data.

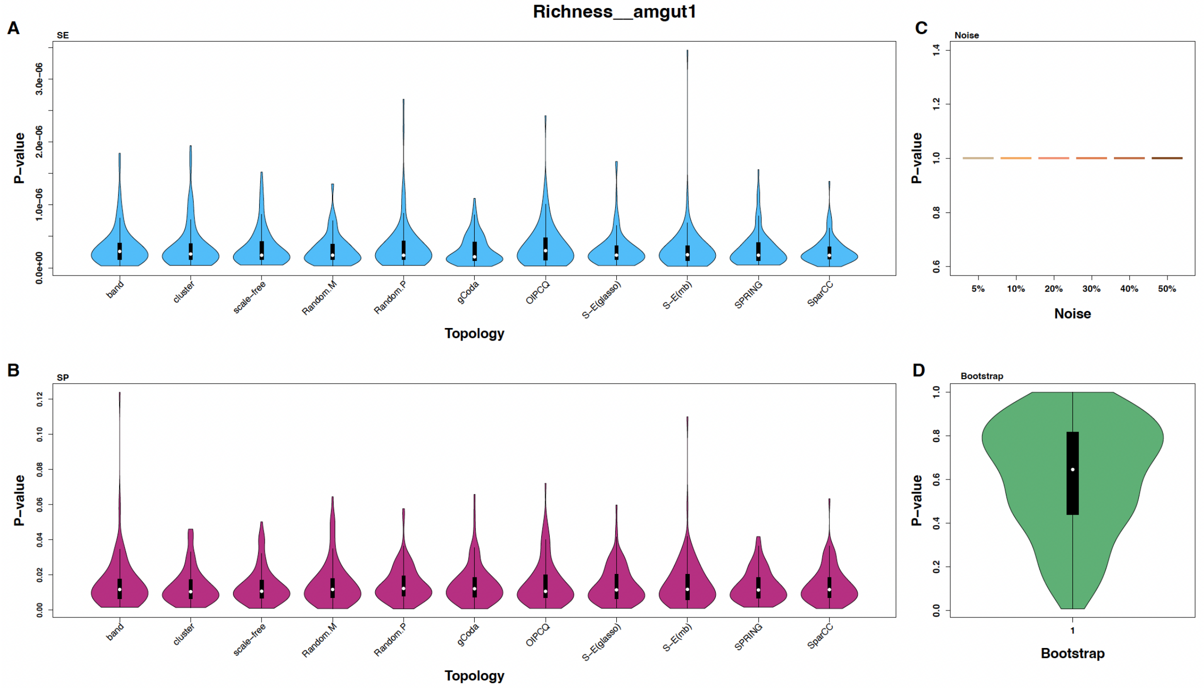


Figure S9: Violin plots showing the distribution of p-values from Wilcoxon tests comparing the Richness index between the `amgut1` dataset and multiple generated datasets. Panels display results for (A) SE-generated synthetic datasets, (B) SP-generated synthetic datasets, (C) noisy datasets with varying noise levels, and (D) bootstrap datasets.

The p-values for all synthetic datasets are below 0.05, indicating that these data significantly differ from the `amgut1` data regarding the Richness index. Furthermore, the synthetic data from the SE method demonstrate a greater difference compared to the SP data. Additionally the noisy and bootstrap datasets, do not exhibit significant differences according to the Wilcoxon test.

The results of the Wilcoxon test for Richness values in the `amgut2`, `GUT`, and `MOMS-PI` datasets are shown in Figures S10, S11, and S12, respectively.

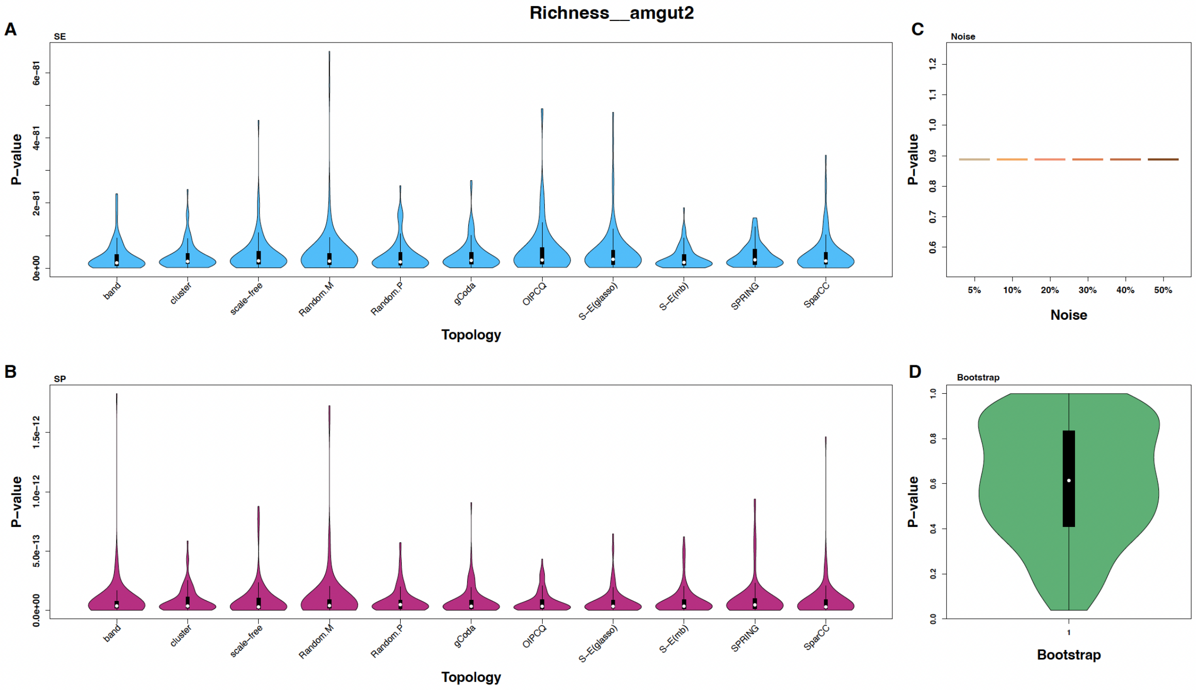


Figure S10: Violin plots showing the distribution of p-values from Wilcoxon tests comparing the Richness index between the `amgut2` dataset and multiple generated datasets. Panels display results for (A) SE-generated synthetic datasets, (B) SP-generated synthetic datasets, (C) noisy datasets with varying noise levels, and (D) bootstrap datasets.

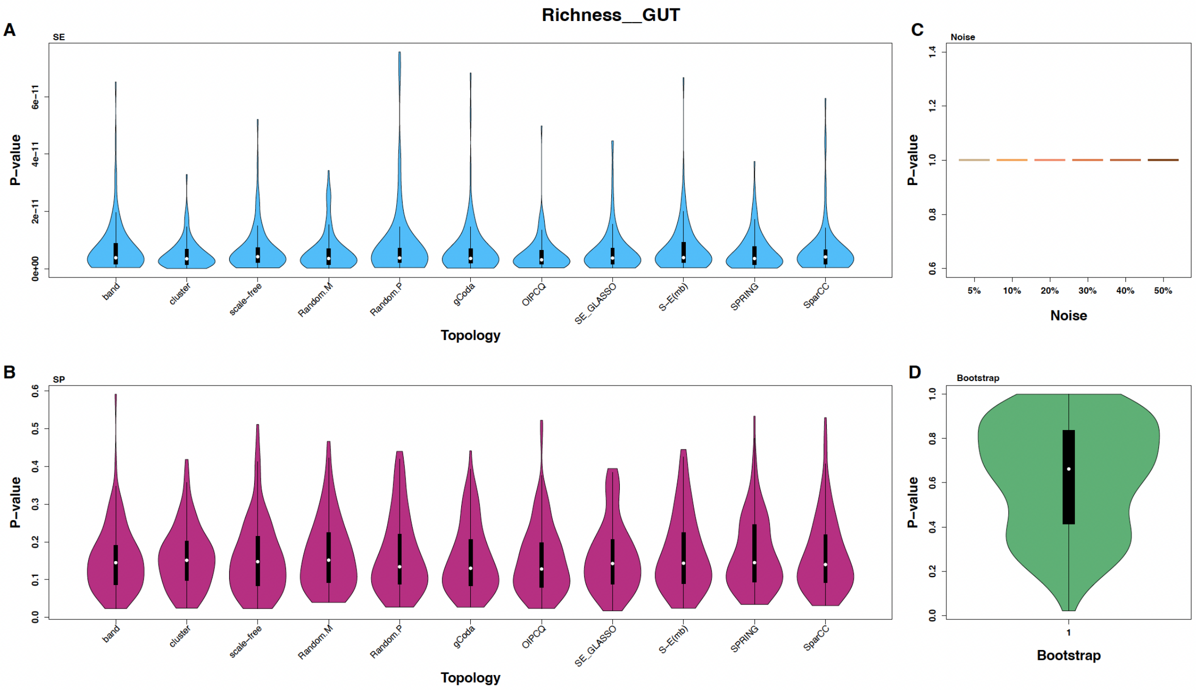


Figure S11: Violin plots showing the distribution of p-values from Wilcoxon tests comparing the Richness index between the `GUT` dataset and multiple generated datasets. Panels display results for (A) SE-generated synthetic datasets, (B) SP-generated synthetic datasets, (C) noisy datasets with varying noise levels, and (D) bootstrap datasets.

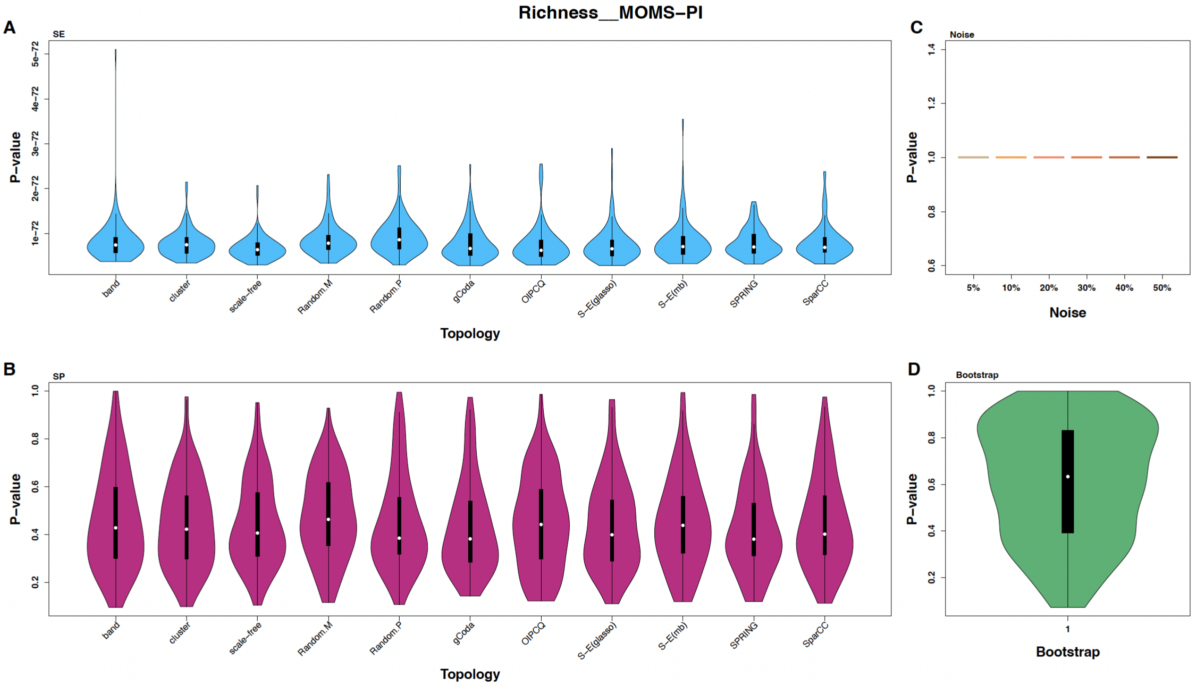


Figure S12: Violin plots showing the distribution of p-values from Wilcoxon tests comparing the Richness index between the `MOMS-PI` dataset and multiple generated datasets. Panels display results for (A) SE-generated synthetic datasets, (B) SP-generated synthetic datasets, (C) noisy datasets with varying noise levels, and (D) bootstrap datasets.

### 2 Comparison of OTU Abundance Distributions Between Real-World and Generated (Synthetic, Noisy, Bootstrap) Datasets

To analyze the distribution of OTUs in the generated data compared to the real dataset, we utilized the Kolmogorov-Smirnov (KS) test. Specifically, we evaluated the distributional differences of each OTU between the real data and each of the other generated datasets using the KS test. A p-value of less than 0.05 from this test indicates a statistically significant difference. For each dataset, we counted the number of OTUs showing a significant difference. Figure S13 displays a violin plot illustrating the percentage of OTUs from each data type that demonstrate a significant difference for the `amgut1` dataset.

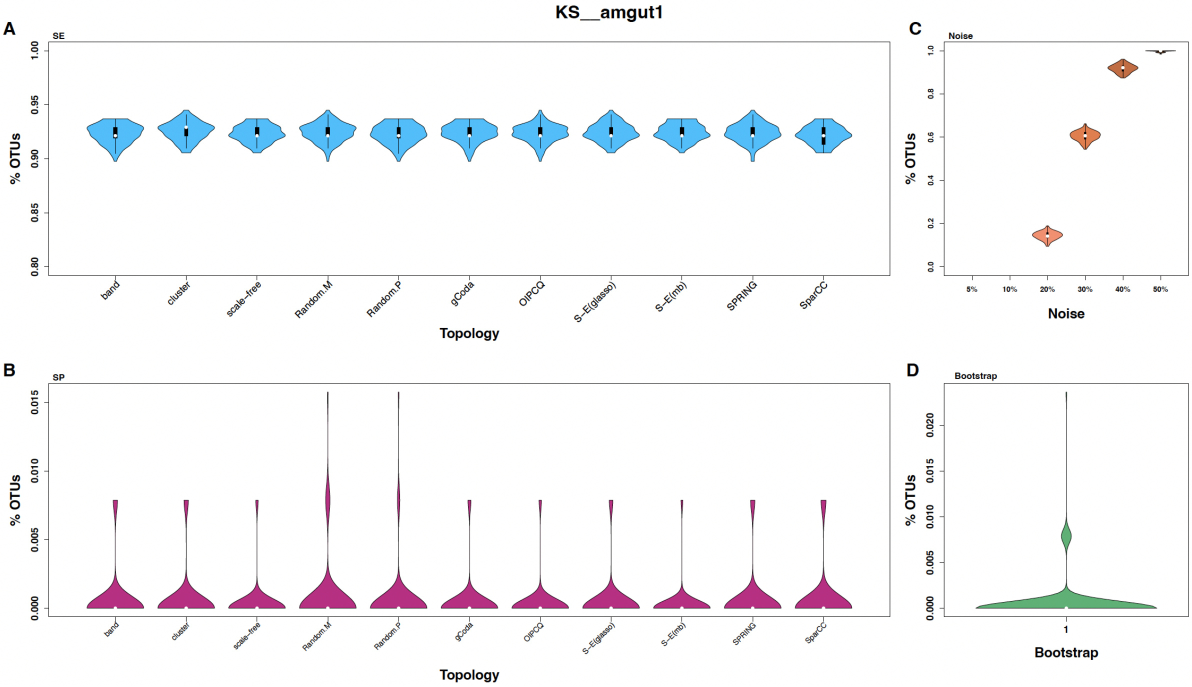


Figure S13: Violin plots illustrating the percentage of OTUs exhibiting significantly different abundance distributions compared to the real `amgut1` dataset, as determined by KS tests ($p < 0.05$). Panels display results for (A) SE-generated synthetic datasets, (B) SP-generated synthetic datasets, (C) noisy datasets with varying noise levels, and (D) bootstrap datasets.

For the `amgut1` dataset, synthetic SE data demonstrate a significant difference in the distribution of OTUs in an average of 92 % of cases, while synthetic SP data reveal almost no significant difference in OTU distributions. Similarly, bootstrapped data also indicate no significant difference in their OTU distributions. Noisy data with up to 10 % noise show no significant difference For noise levels of 20 %, 30 %, 40 %, and 50 %, there are significant differences between the OTU distributions in 14 %, 60 %, 92 %, and 100 % of cases, respectively.

The results of the KS test for the `amgut2`, `GUT`, and `MOMS-PI` datasets are presented in Figures S14, S15, and S16, respectively.

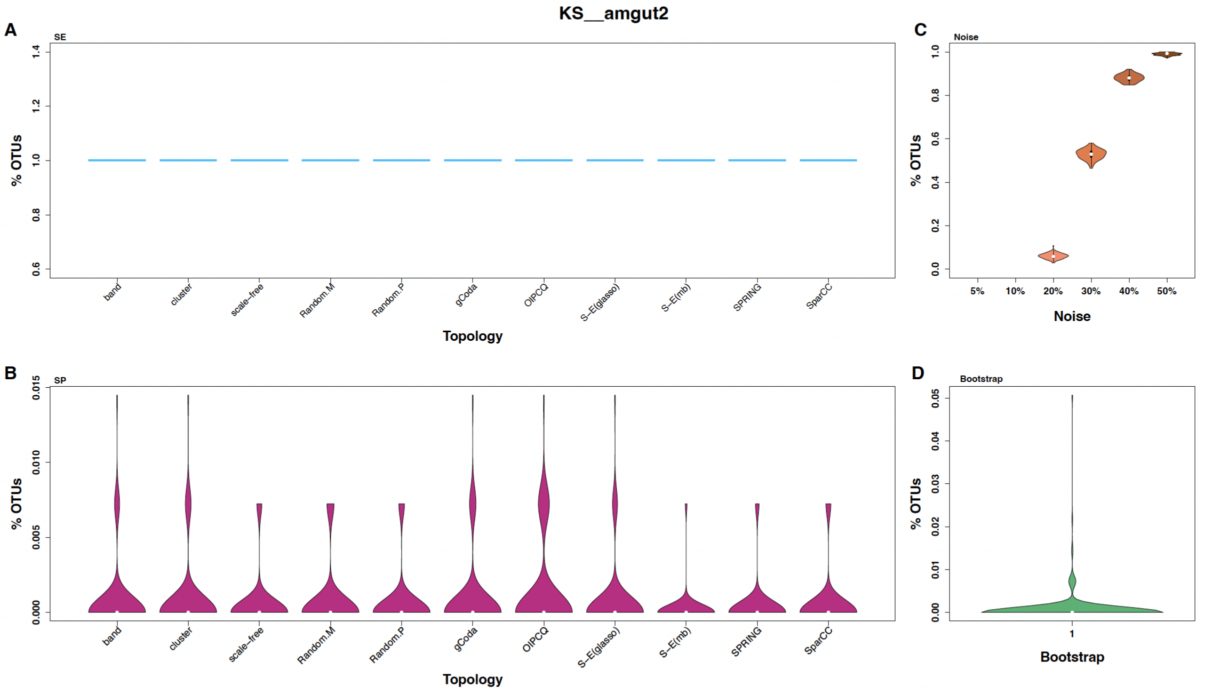


Figure S14: Violin plots illustrating the percentage of OTUs exhibiting significantly different abundance distributions compared to the real `amgut2` dataset, as determined by KS tests ($p < 0.05$). Panels display results for (A) SE-generated synthetic datasets, (B) SP-generated synthetic datasets, (C) noisy datasets with varying noise levels, and (D) bootstrap datasets.

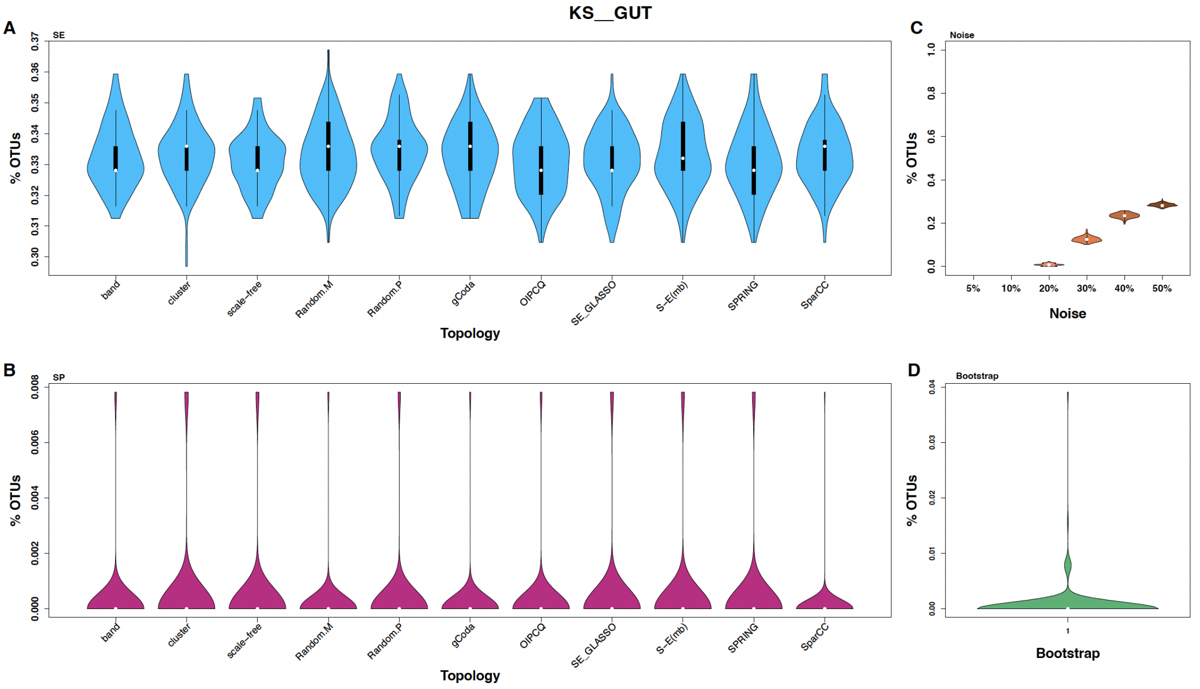


Figure S15: Violin plots illustrating the percentage of OTUs exhibiting significantly different abundance distributions compared to the real `GUT` dataset, as determined by KS tests ($p < 0.05$). Panels display results for (A) SE-generated synthetic datasets, (B) SP-generated synthetic datasets, (C) noisy datasets with varying noise levels, and (D) bootstrap datasets.

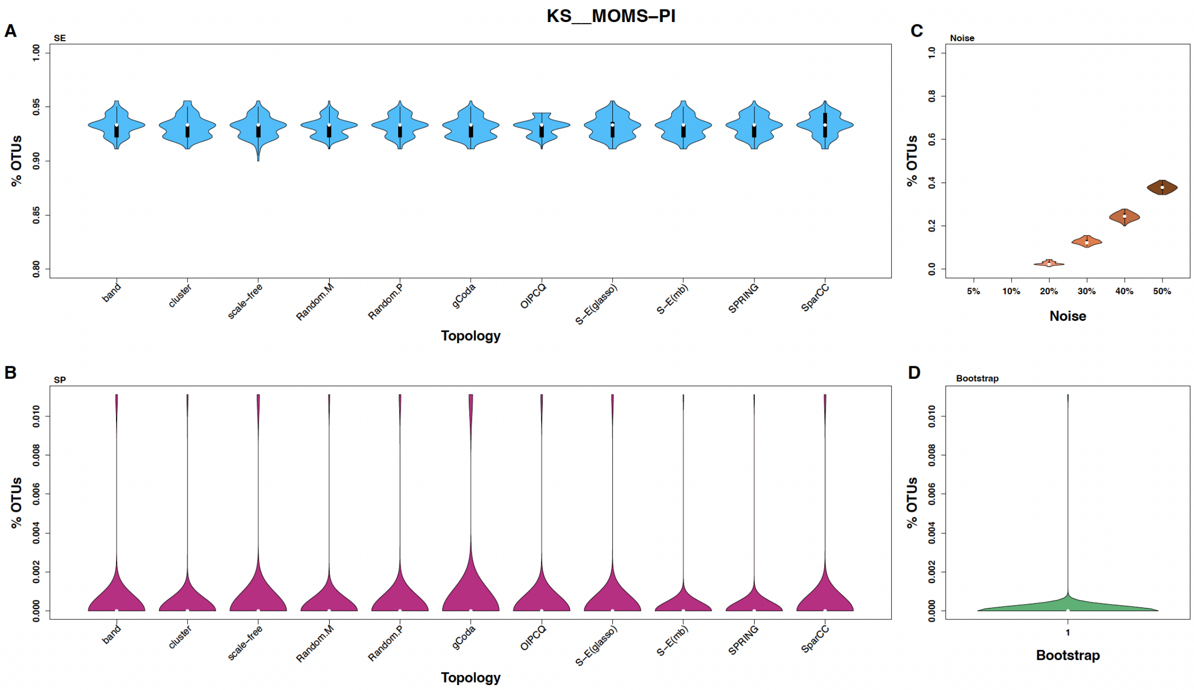


Figure S16: Violin plots illustrating the percentage of OTUs exhibiting significantly different abundance distributions compared to the real `MOMS-PI` dataset, as determined by KS tests ($p < 0.05$). Panels display results for (A) SE-generated synthetic datasets, (B) SP-generated synthetic datasets, (C) noisy datasets with varying noise levels, and (D) bootstrap datasets.

### 3 Algorithms Performance in Microbial Network Inference Using Generated Datasets

#### 3.1 Precision and Recall for amgut1

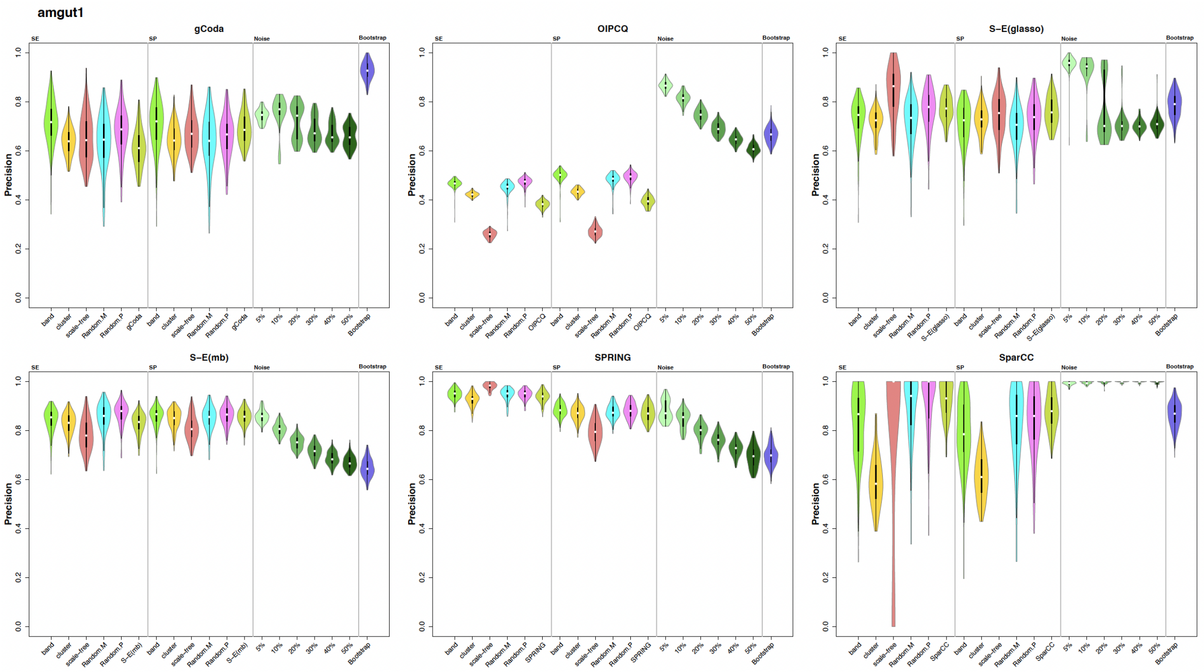


Figure S17: Violin plots showing the distribution of Precisions for six network inference algorithms applied to the `amgut1` dataset across synthetic (SE and SP), noisy, and bootstrap data.

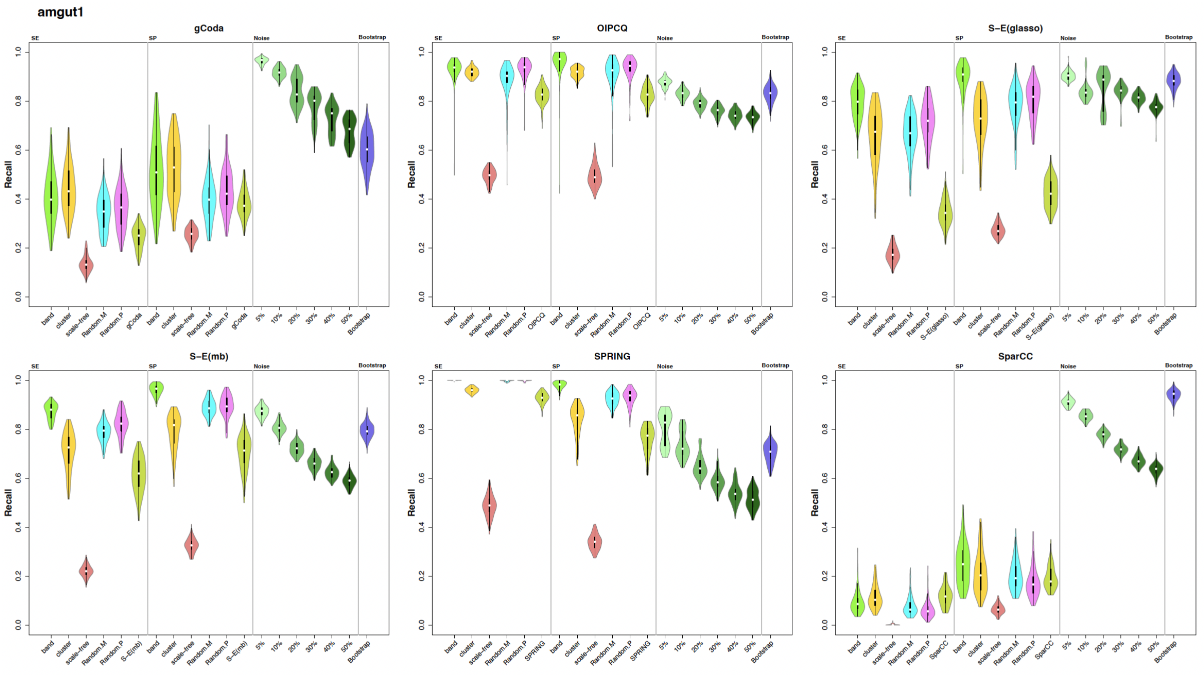


Figure S18: Violin plots showing the distribution of Recalls for six network inference algorithms applied to the `amgut1` dataset across synthetic (SE and SP), noisy, and bootstrap data.

#### 3.2 Precision, Recall and F-score for amgut2

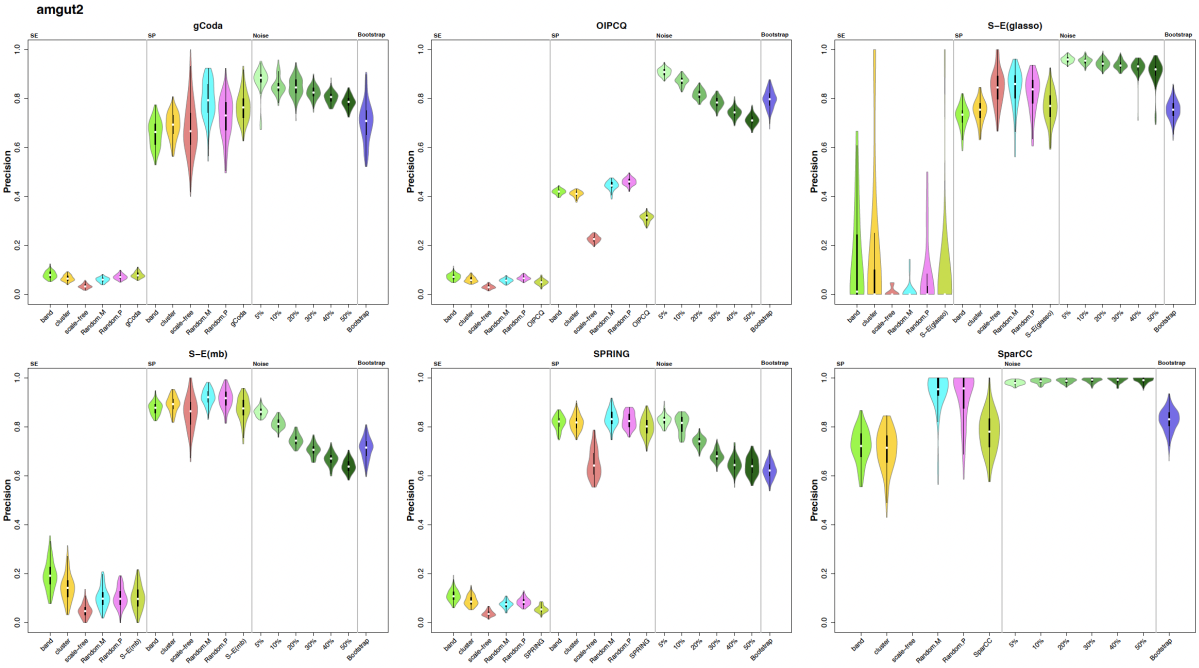


Figure S19: Violin plots showing the distribution of Precisions for six network inference algorithms applied to the `amgut2` dataset across synthetic (SE and SP), noisy, and bootstrap data.

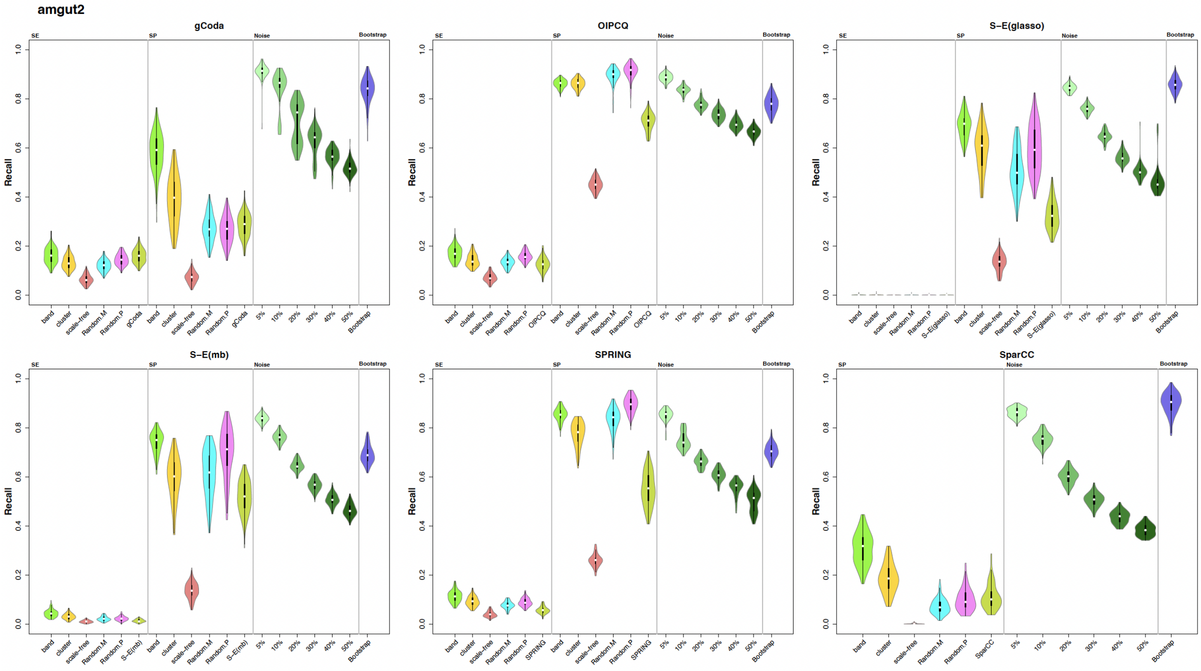


Figure S20: Violin plots showing the distribution of Recalls for six network inference algorithms applied to the `amgut2` dataset across synthetic (SE and SP), noisy, and bootstrap data.

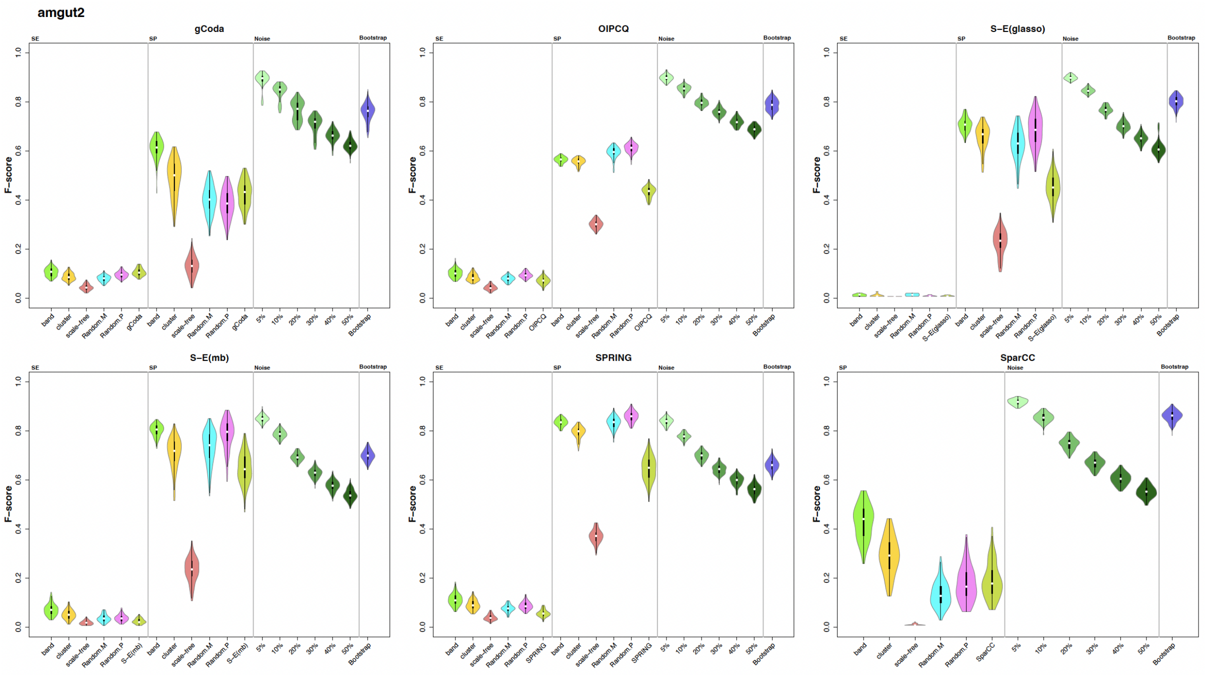


Figure S21: Violin plots showing the distribution of F-scores for six network inference algorithms applied to the `amgut2` dataset across synthetic (SE and SP), noisy, and bootstrap data.

#### 3.3 Precision, Recall and F-score for GUT

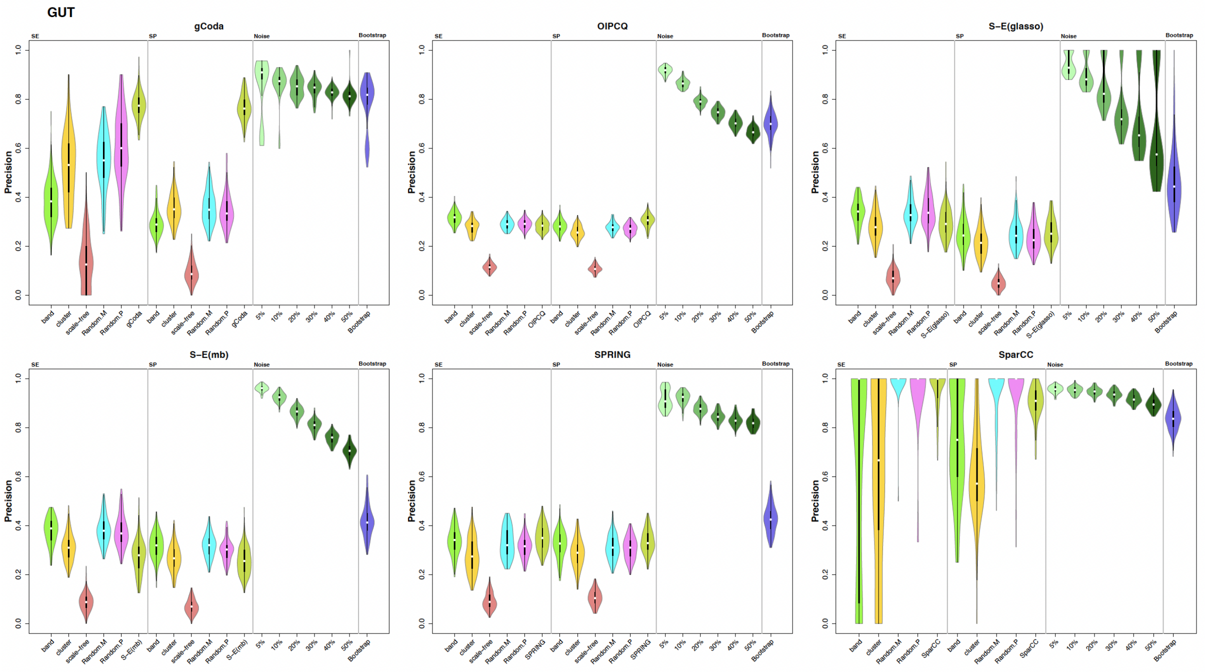


Figure S22: Violin plots showing the distribution of Precisions for six network inference algorithms applied to the `GUT` dataset across synthetic (SE and SP), noisy, and bootstrap data.

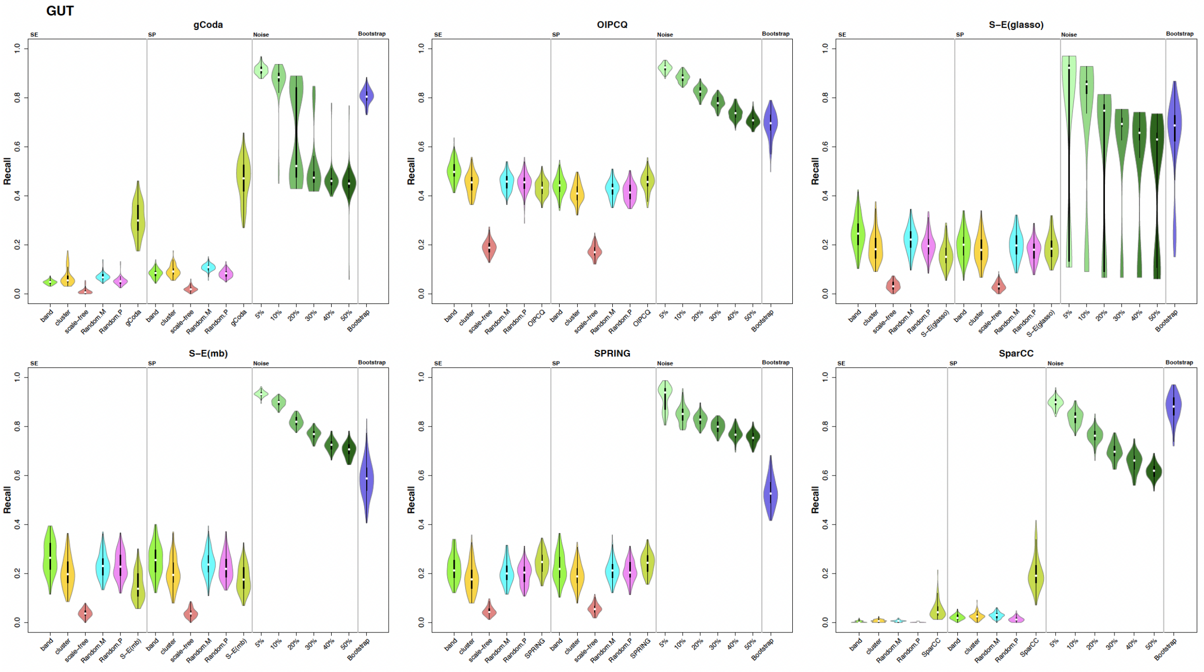


Figure S23: Violin plots showing the distribution of Recalls for six network inference algorithms applied to the GUT dataset across synthetic (SE and SP), noisy, and bootstrap data.

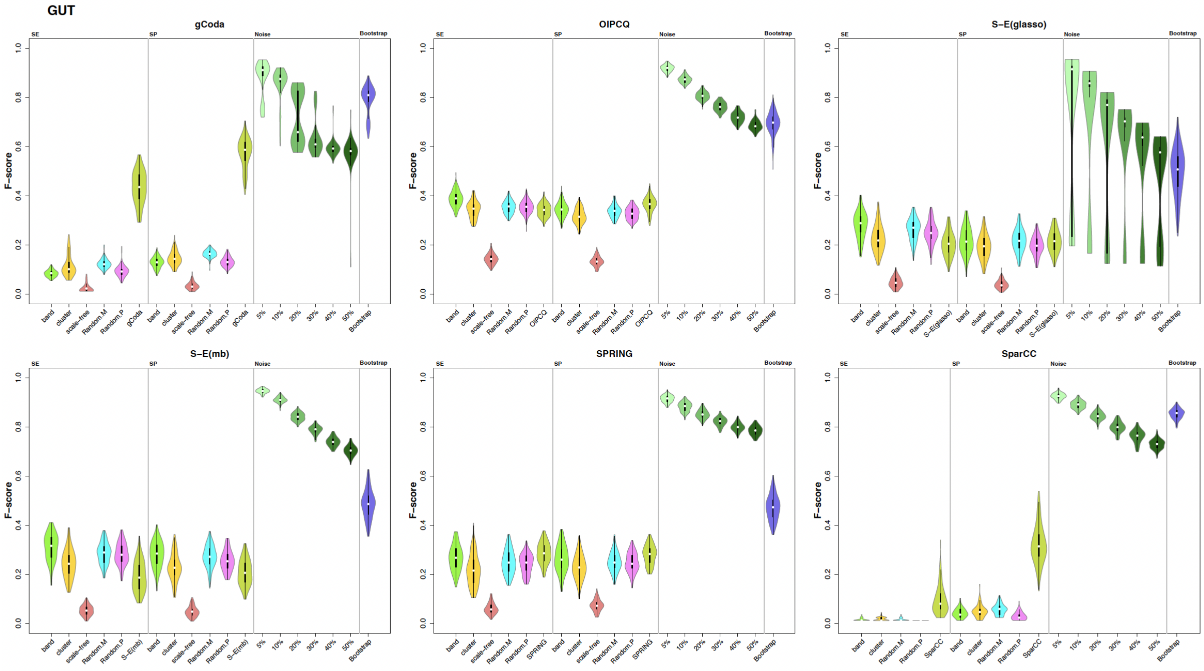


Figure S24: Violin plots showing the distribution of F-scores for six network inference algorithms applied to the GUT dataset across synthetic (SE and SP), noisy, and bootstrap data.

#### 3.4 Precision, Recall and F-score for MOMS-PI

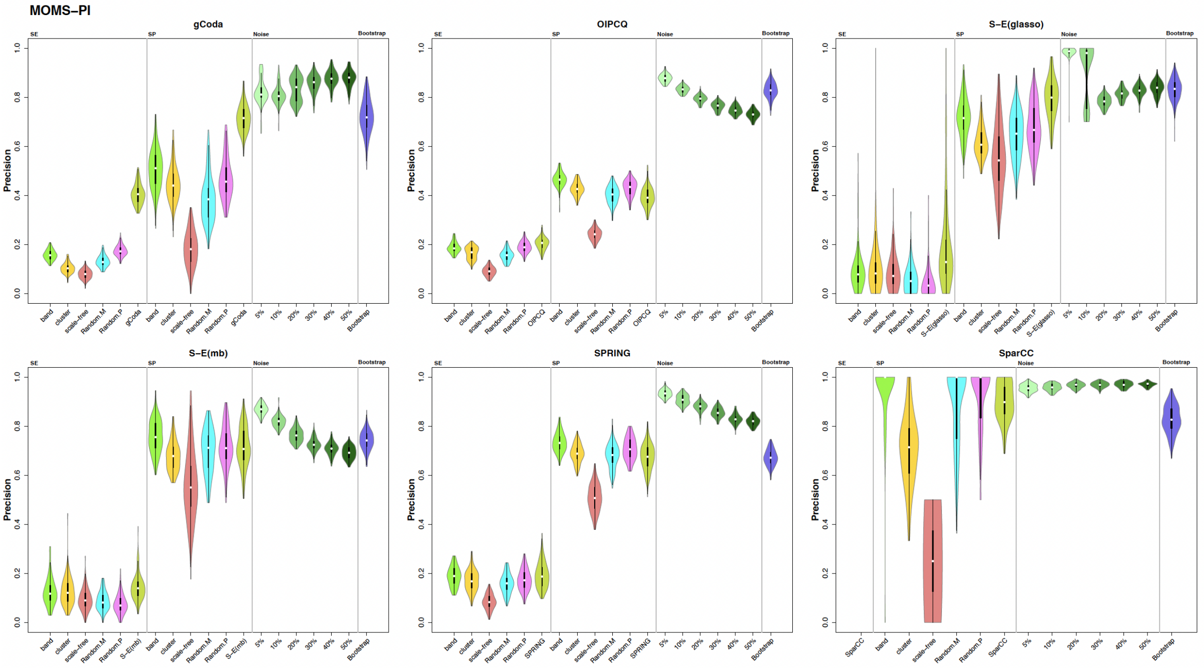


Figure S25: Violin plots showing the distribution of Precisions for six network inference algorithms applied to the MOMS-PI dataset across synthetic (SE and SP), noisy, and bootstrap data.

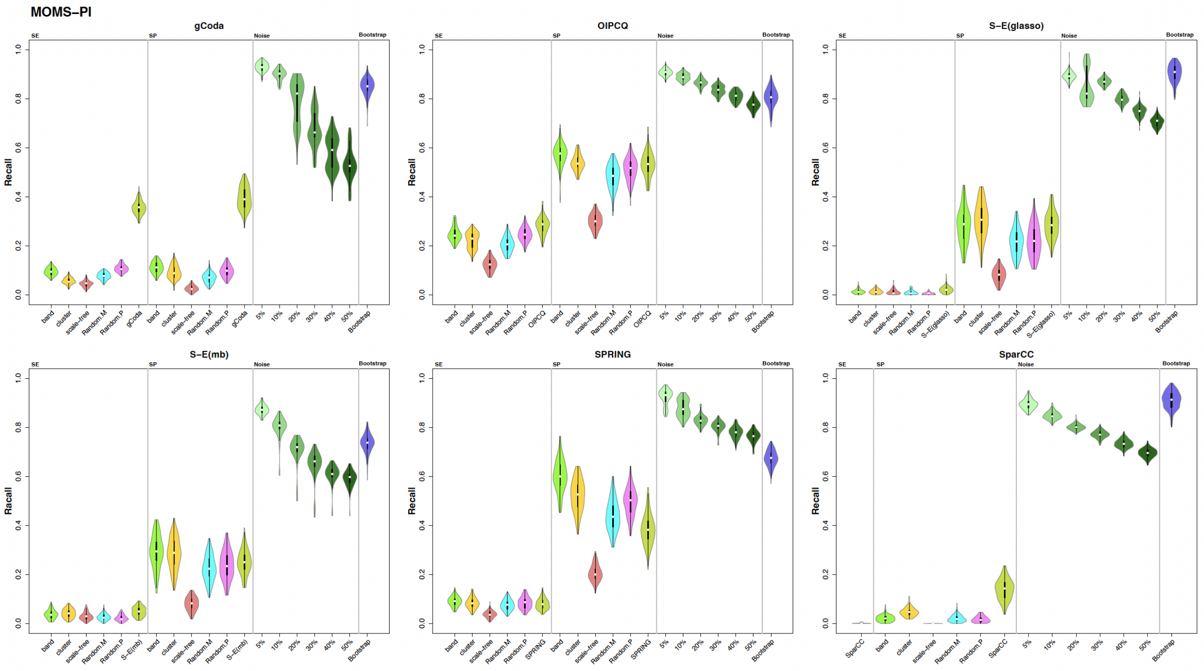


Figure S26: Violin plots showing the distribution of Recalls for six network inference algorithms applied to the MOMS-PI dataset across synthetic (SE and SP), noisy, and bootstrap data.

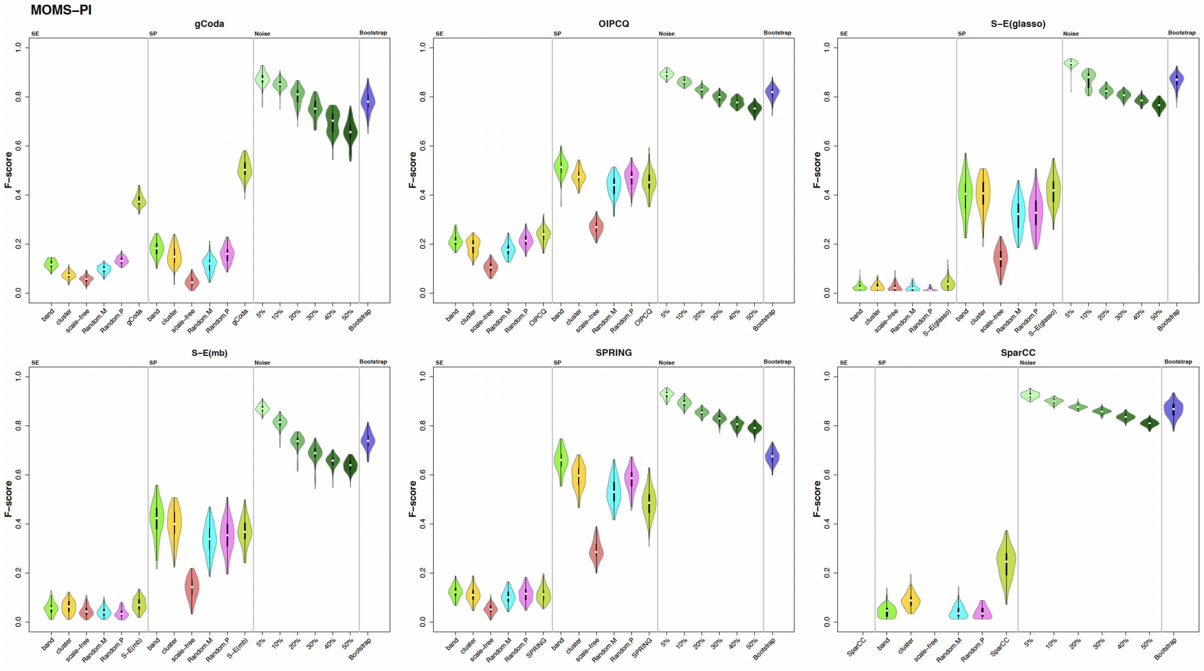


Figure S27: Violin plots showing the distribution of F-scores for six network inference algorithms applied to the MOMS-PI dataset across synthetic (SE and SP), noisy, and bootstrap data.

### 4 Pairwise Statistical Comparisons Among Generated Data Results

This section presents a series of visualizations that assess the consistency and variability of the results across different data variants. The heatmap illustrates the pairwise statistical differences among the outcomes from various generated data sets, providing insights into the homogeneity and stability of the data generation process.

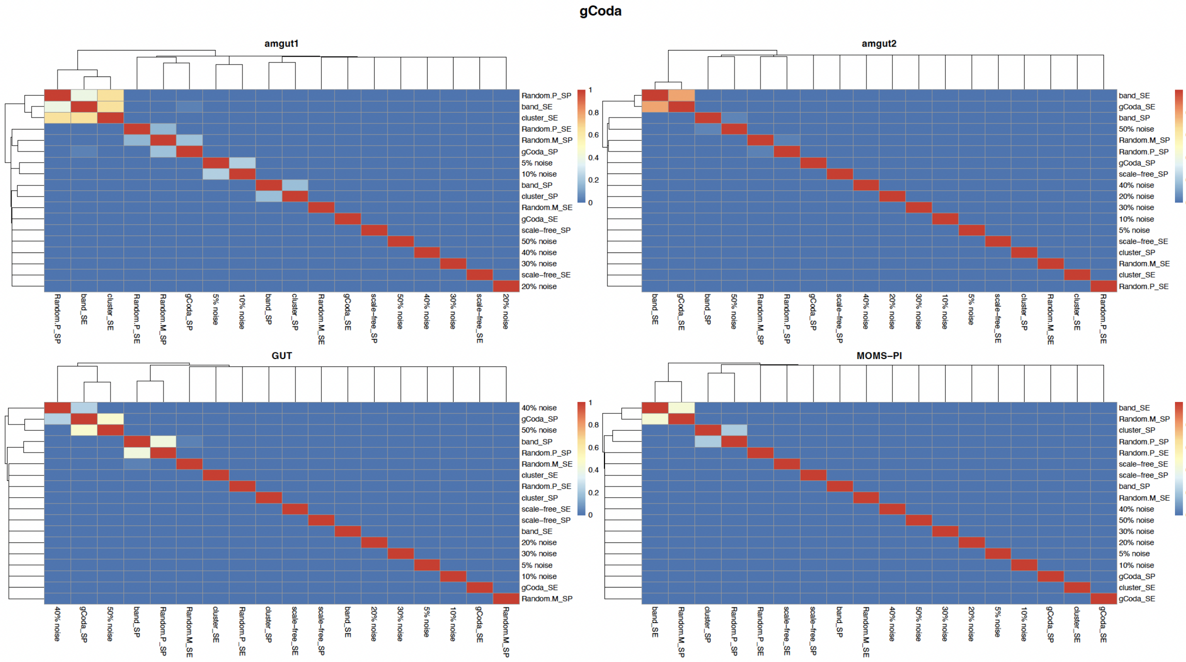


Figure S28: Heatmap showing pairwise statistical comparisons of results from different data variants. This visualization displays the results of t-tests or Wilcoxon tests, where $p - value > 0.05$ indicates no significant difference between pairs. It assesses the consistency and stability of outcomes across data generation methods.

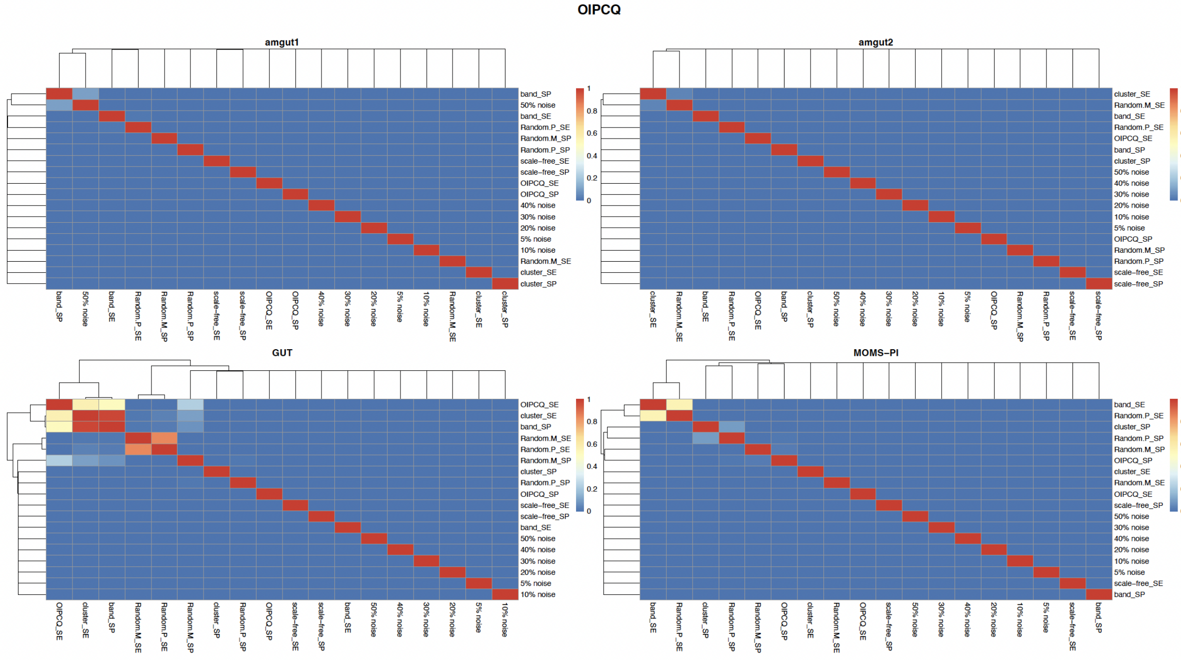


Figure S29: Heatmap showing pairwise statistical comparisons of results from different data variants. This visualization displays the results of t-tests or Wilcoxon tests, where $p - value > 0.05$ indicates no significant difference between pairs. It assesses the consistency and stability of outcomes across data generation methods.

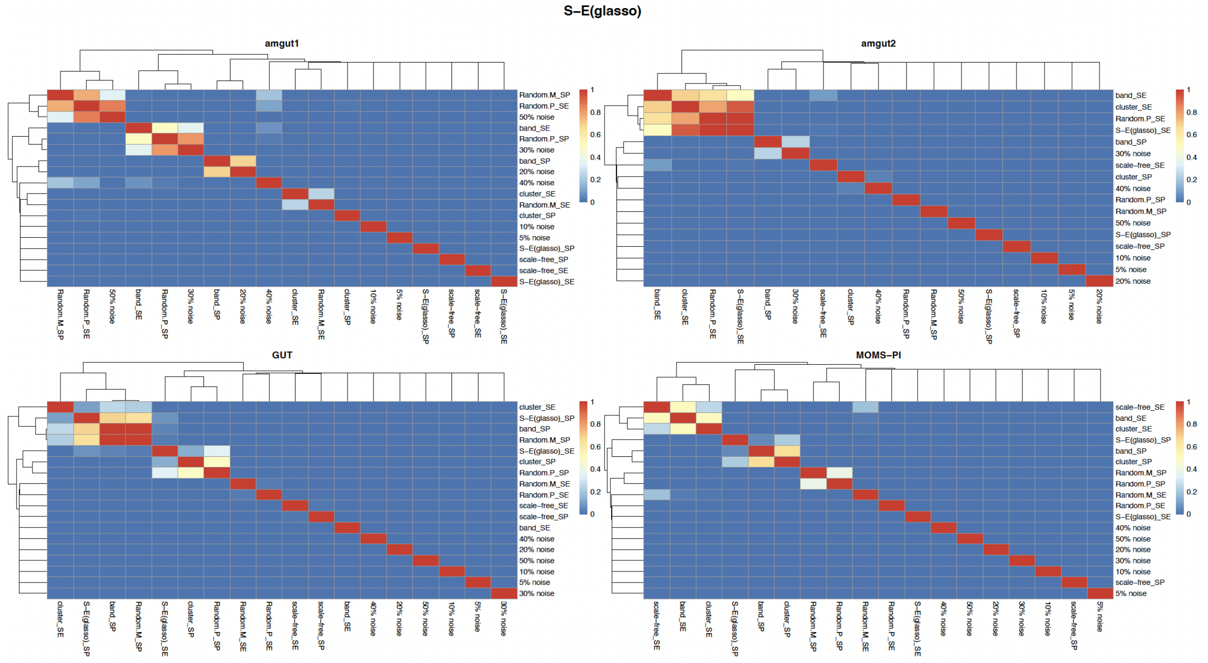


Figure S30: Heatmap showing pairwise statistical comparisons of results from different data variants. This visualization displays the results of t-tests or Wilcoxon tests, where $p - value > 0.05$ indicates no significant difference between pairs. It assesses the consistency and stability of outcomes across data generation methods.

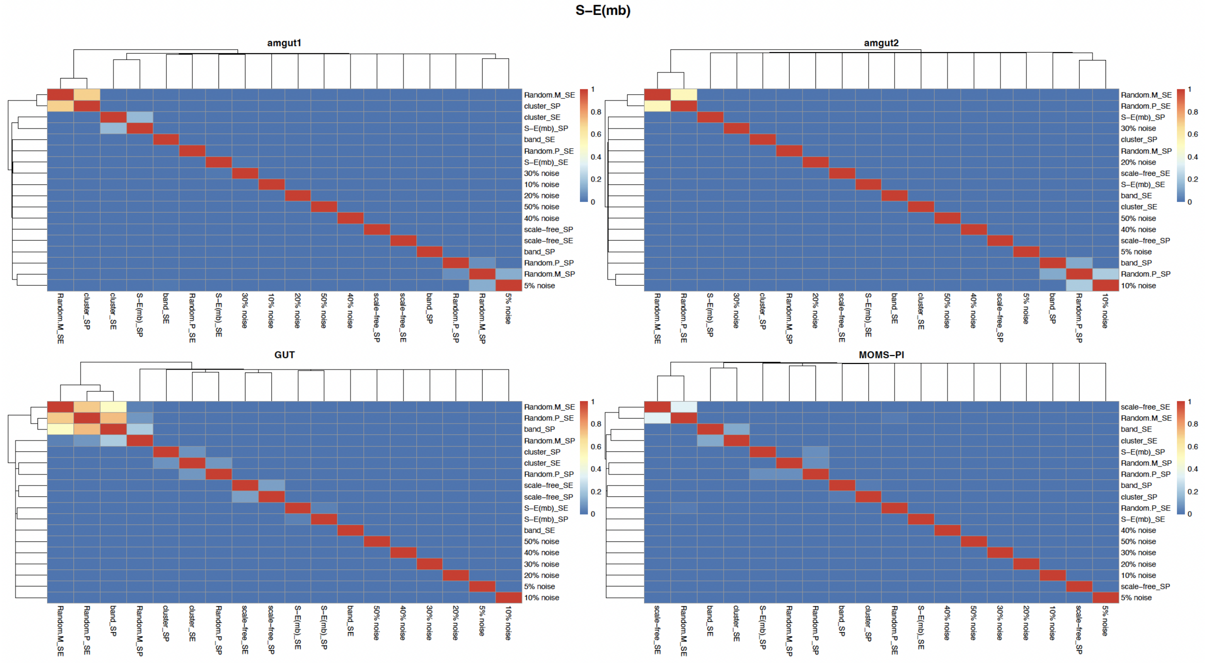


Figure S31: Heatmap showing pairwise statistical comparisons of results from different data variants. This visualization displays the results of t-tests or Wilcoxon tests, where $p - value > 0.05$ indicates no significant difference between pairs. It assesses the consistency and stability of outcomes across data generation methods.

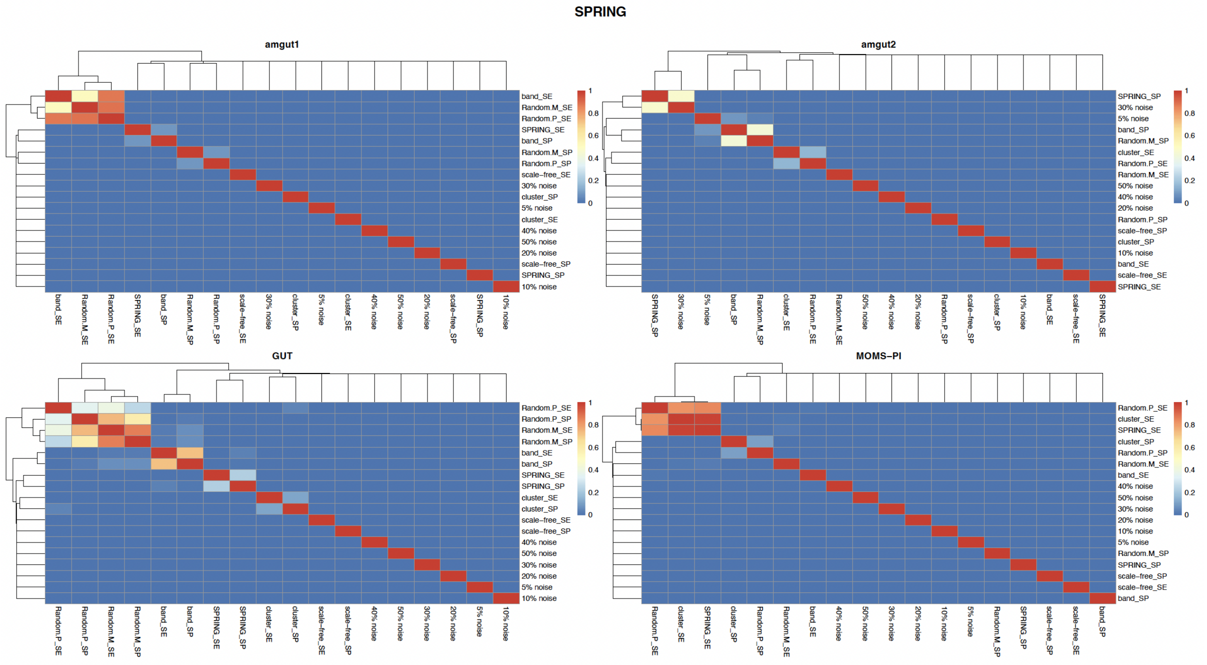


Figure S32: Heatmap showing pairwise statistical comparisons of results from different data variants. This visualization displays the results of t-tests or Wilcoxon tests, where $p - value > 0.05$ indicates no significant difference between pairs. It assesses the consistency and stability of outcomes across data generation methods.

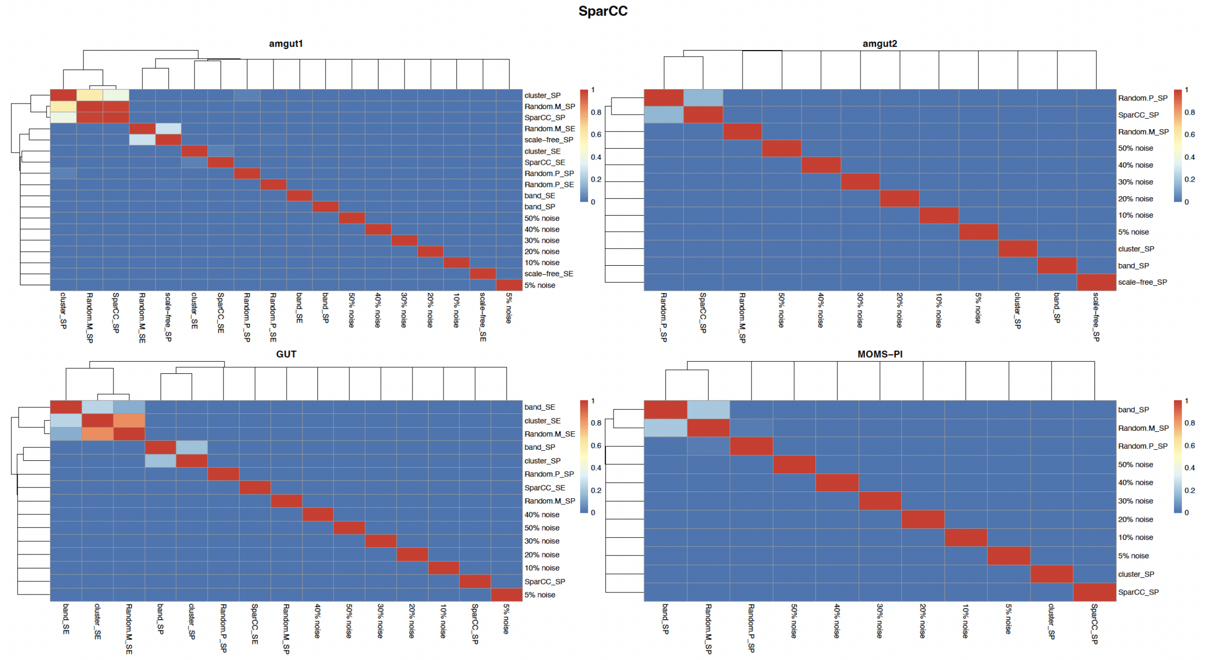


Figure S33: Heatmap showing pairwise statistical comparisons of results from different data variants. This visualization displays the results of t-tests or Wilcoxon tests, where $p - value > 0.05$ indicates no significant difference between pairs. It assesses the consistency and stability of outcomes across data generation methods.

### 5 Comparison of Algorithms Performance Across All Generated Datasets for Real-World Data

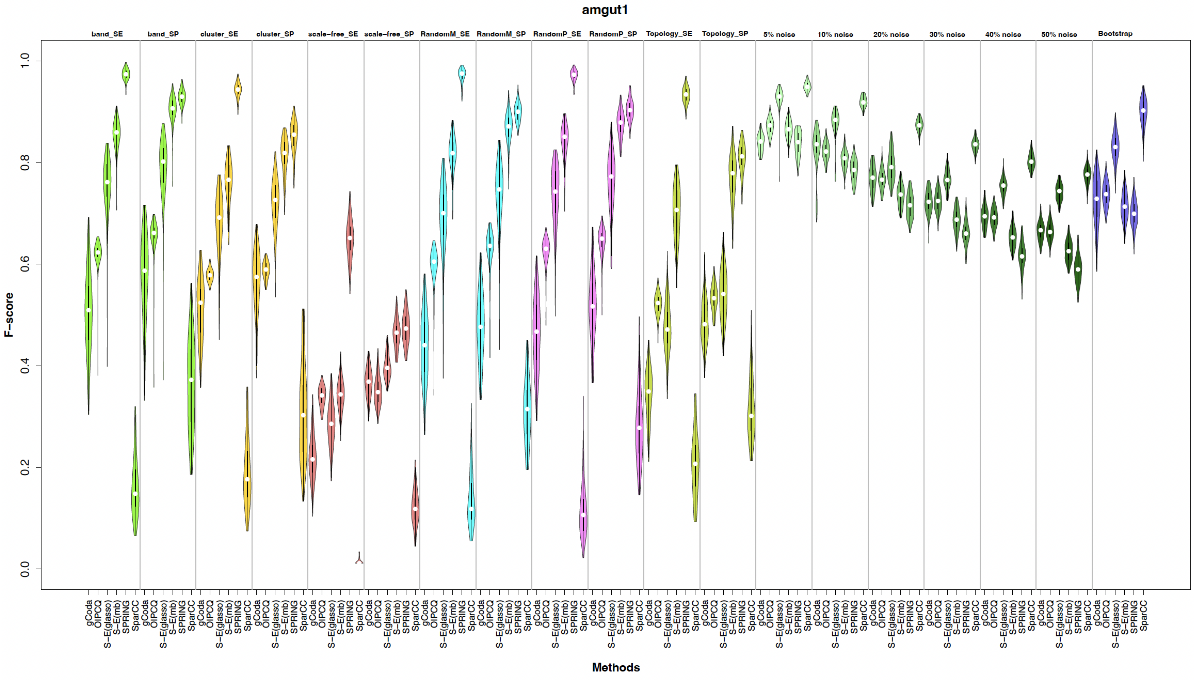


Figure S34: Violin plots comparing F-score distributions of all six algorithms across all generated datasets from the `amgut1`. *"band_SE"* represents datasets generated from the band topology using the SE method; *"Topology_SE"* denotes datasets generated based on each algorithm's inferred topology using the SE method. These comparisons highlight algorithm robustness and performance consistency across diverse data generation strategies.

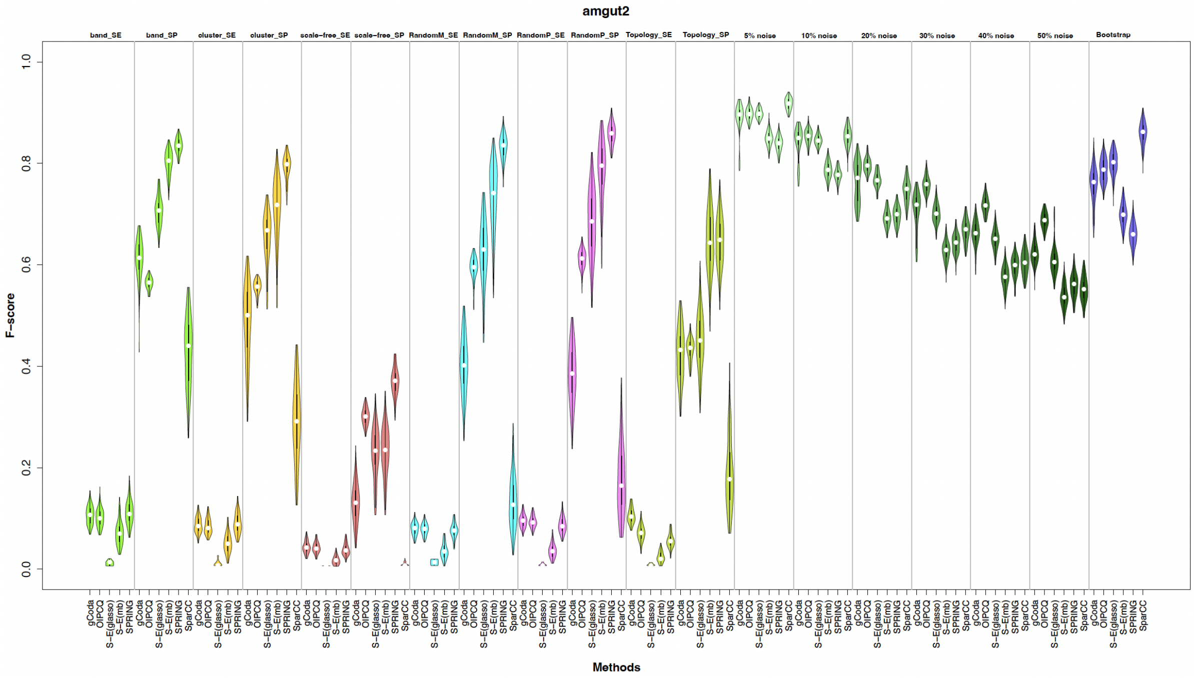


Figure S35: Violin plots comparing F-score distributions of all six algorithms across all generated datasets from the `amgut2`. *"band_SE"* represents datasets generated from the band topology using the SE method; *"Topology_SE"* denotes datasets generated based on each algorithm's inferred topology using the SE method. These comparisons highlight algorithm robustness and performance consistency across diverse data generation strategies.

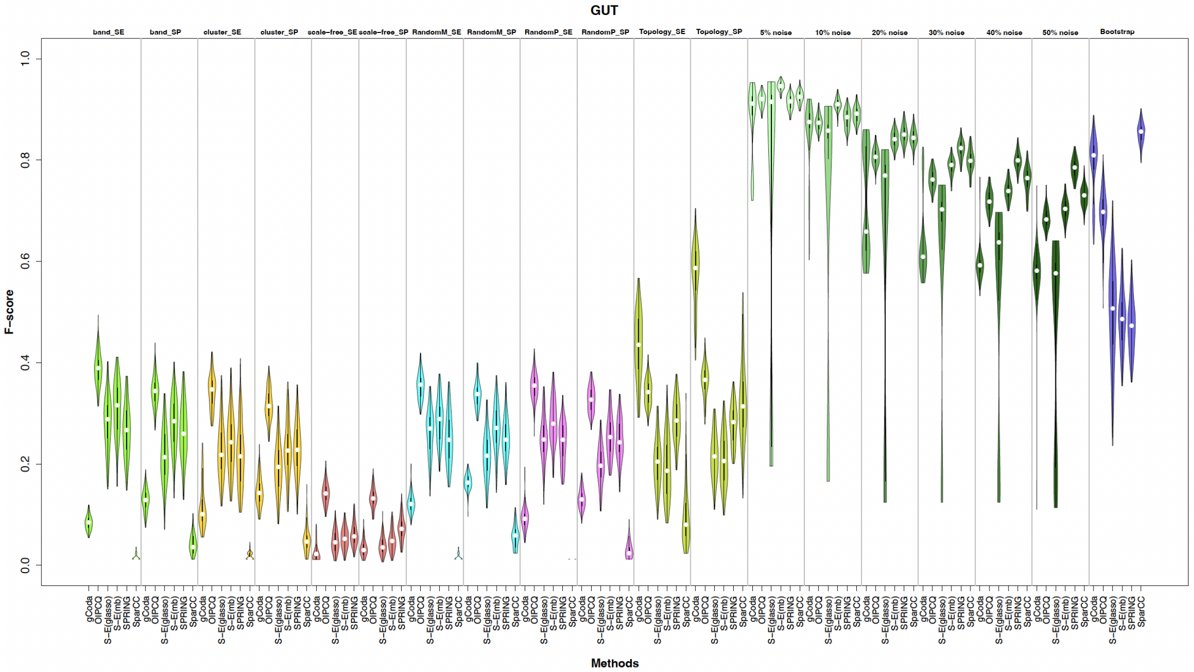


Figure S36: Violin plots comparing F-score distributions of all six algorithms across all generated datasets from the `GUT`. *"band_SE"* represents datasets generated from the band topology using the SE method; *"Topology_SE"* denotes datasets generated based on each algorithm's inferred topology using the SE method. These comparisons highlight algorithm robustness and performance consistency across diverse data generation strategies.

### 6 Algorithm Performance Under Varying Noise Levels

In this section, we compare algorithms performance as noise intensity increases.

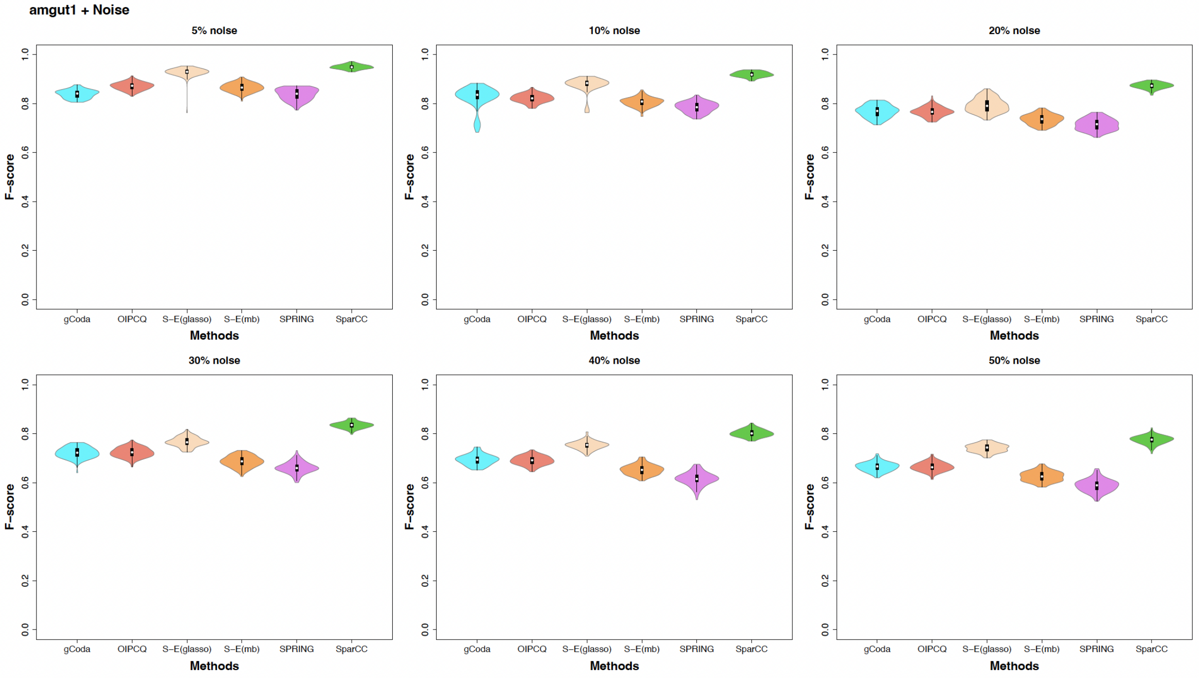


Figure S37: Violin plots comparing F-scores of different algorithms across varying noise levels in the `amgut1` dataset.

In `amgut1` for 5 % noise, the results of the gCoda and SPRING algorithms do not show a significant difference.

For 20 %, 30 %, 40 %, and 50 % noise levels, the results of the gCoda and OIPCQ algorithms also do not exhibit statistically significant differences.

For all noise levels in the `amgut1` dataset, the SparCC algorithm provides the best results with the least variance.

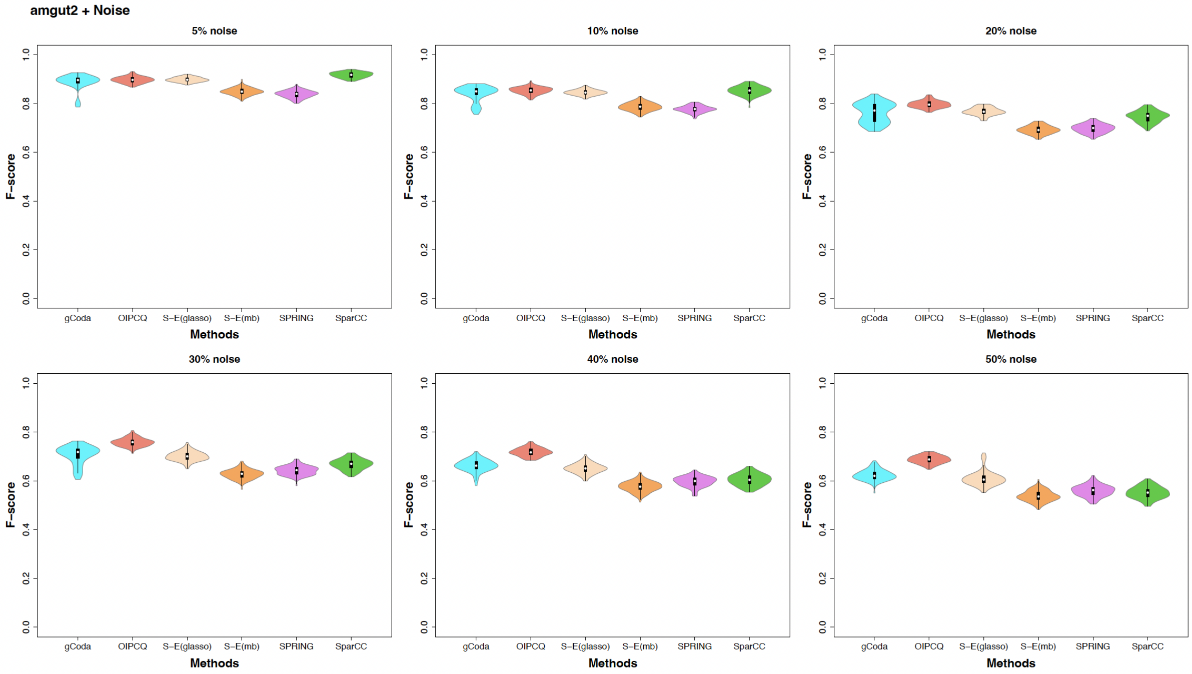


Figure S38: Violin plots comparing F-scores of different algorithms across varying noise levels in the `amgut2` dataset.

In `amgut2` for 5 % noise, the results of the gCoda algorithm do not show a statistically significant difference compared to the OIPCQ and S-E(glasso) algorithms. Furthermore, the OIPCQ and S-E(glasso) algorithms also do not exhibit a statistically significant difference.

For 10 % noise, the results of the gCoda algorithm do not show a statistically significant difference compared to the OIPCQ, S-E(glasso), and SparCC algorithms. Additionally, the OIPCQ and SparCC algorithms do not exhibit a statistically significant difference.

For 20 % noise, the results of the gCoda and S-E(glasso) algorithms do not show a statistically significant difference.

For 5 % and 10 % noise levels, the best result for the `amgut2` dataset is obtained by the SparCC algorithm. For higher noise levels, the OIPCQ algorithm achieves the best result.

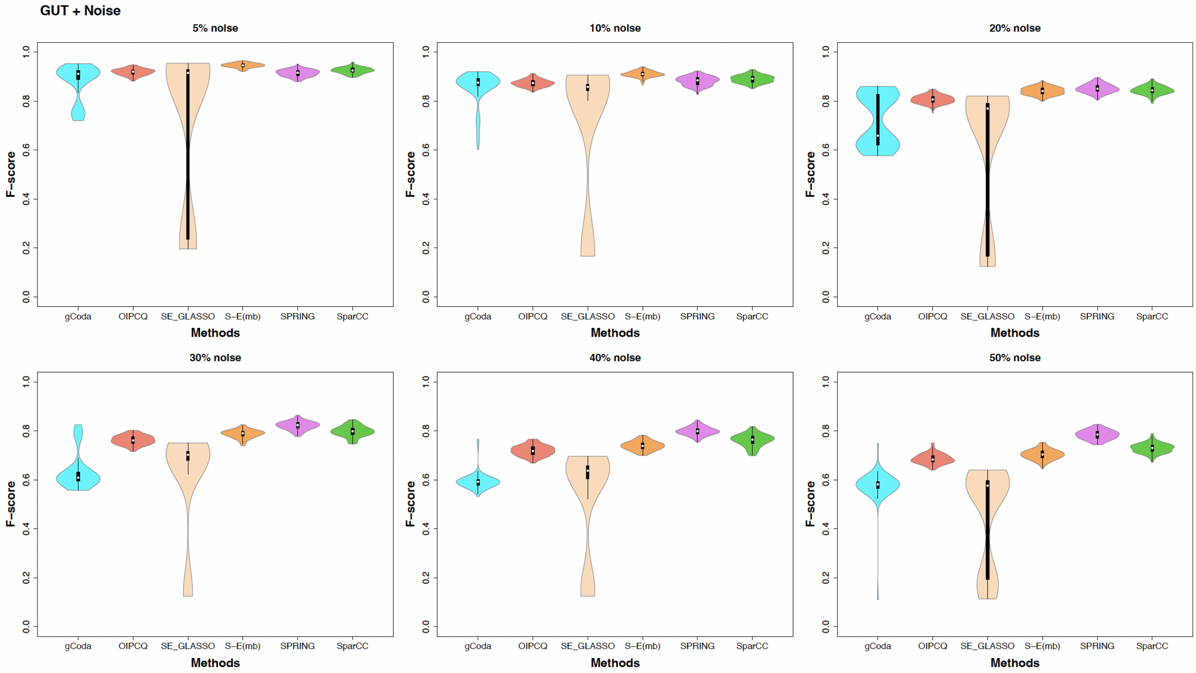


Figure S39: Violin plots comparing F-scores of different algorithms across varying noise levels in the GUT dataset.

In GUT for 5 % noise, the results of the gCoda algorithm do not show a statistically significant difference compared to the S-E(glasso) and SPRING algorithms. Furthermore, the OIPCQ and S-E(glasso) algorithms, as well as S-E(glasso) and SPRING, do not exhibit statistically significant differences.

For 10 % noise, the results of the gCoda algorithm do not show a statistically significant difference compared to the OIPCQ and SPRING algorithms.

For 20 % noise, the results of the S-E(mb) and SparCC algorithms do not show a statistically significant difference.

For noise levels of 5 % and 10 %, the S-E(mb) method yields the best results. For higher noise levels, the best results are obtained from the SPRING algorithm.

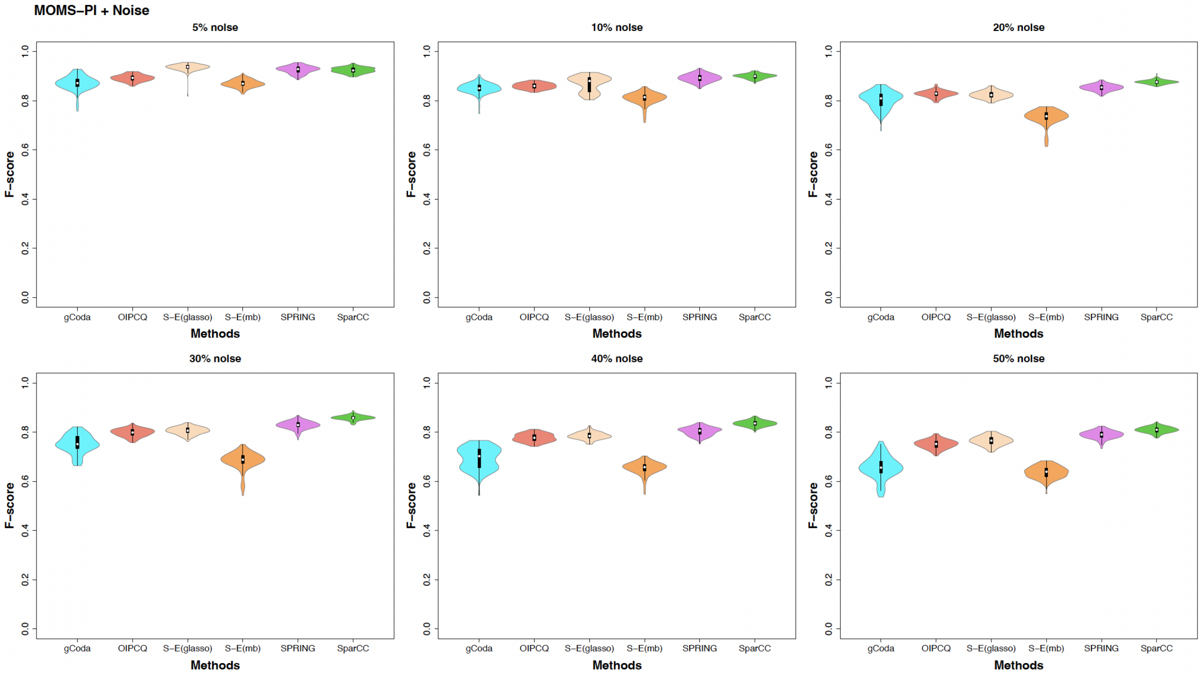


Figure S40: Violin plots comparing F-scores of different algorithms across varying noise levels in the MOMS-PI dataset.

In MOMS-PI for 5 % noise, the results of the gCoda and S-E(mb) algorithms do not show a statistically significant difference. Similarly, the SPRING and SparCC algorithms also lack a statistically significant difference.

For 20 % noise, the results of the OIPCQ and S-E(glasso) algorithms do not show a statistically significant difference.

For 5 % noise, the best result is obtained from the S-E(glasso) algorithm. For higher noise levels, the best result is obtained from the SparCC algorithm.
